# Genomic, genetic and phylogenetic evidence for a new falcon species using chromosome-level genome assembly of the gyrfalcon and population genomics

**DOI:** 10.1101/2023.02.12.525808

**Authors:** Farooq Omar Al-Ajli, Giulio Formenti, Olivier Fedrigo, Alan Tracey, Ying Sims, Kerstin Howe, Ikdam M. Al-Karkhi, Asmaa Ali Althani, Erich D. Jarvis, Sadequr Rahman, Qasim Ayub

## Abstract

The taxonomic classification of a falcon population found in the Altai region in Asia has been heavily debated for two centuries and previous studies have been inconclusive, hindering a more informed conservation approach. Here, we generated a chromosome-level gyrfalcon reference genome using the Vertebrate Genomes Project (VGP) assembly pipeline. Using whole genome sequences of 44 falcons from different species and populations, including “Altai” falcons, we analyzed their population structure, admixture patterns and demographic history. We find that the Altai falcons are genomically mosaic of saker and gyrfalcon ancestries, and carry distinct W- and mitochondrial-haplotypes that cluster with the lanner falcon. The Altai haplotype diverged 422,000 years ago from the ancestor of sakers and gyrfalcons, both of which, in turn, split 109,000 years ago. The Altai W chromosome includes coding variants that may influence important structural, behavioral and reproductive traits. These findings support the designation of Altai falcons as a distinct falcon species (*Falco altaicus*).

## Introduction: The falcons of the Altai region as a 200-year-old taxonomic puzzle

Falcons from the Altai Mountain regions in Central and East Asia have historically been referred to as *Falco altaicus* and have been the focus of a more than 200-year-old debate (*1, 2*). Their habitat in the Altai region is a wintering zone for both gyrfalcon (*F. rusticolus*) and saker (*F. cherrug*) falcons, which has contributed to the speculation that the Altai region falcons could represent a natural interspecific hybrid population (*3, 4*). In Mongolia, this falcon population is restricted to western parts of the country including the Altai mountain range (*5*). The currently accepted taxonomical classification of these falcons is either a saker population or a saker subspecies, with its distinctive dark morphology interpreted as a population-specific morph (*4, 6*). Indeed, morphologically some Altai region falcons resemble an intermediate form between sakers and gyrfalcons. Despite the significance of resolving their status for the conservation of their populations, some researchers consider them as natural interspecific gyrfalcon-saker hybrids merely based on their resemblance to sakers and/or gyrfalcons (*4, 7*). Yet, some biologists and taxonomists consider Altai region falcons as a separate species, based on a different set of criteria, including its ecological niche (foraging, breeding, and wintering), as it occupies highly elevated mountains, which sakers usually avoid (*2, 7*–*9*). Another point of view treats these falcons as a subspecies of either the gyrfalcon or saker (i.e. *F. cherrug/rusticolus altaicus*) (*10*). On the hybrid hypothesis, it was suggested that Altai region falcons are a natural hybrid of first or subsequent generations between the gyrfalcons and sakers, with the saker genes predominating (*3, 4*). Others speculated that these falcons are indeed hybrids, but of falconry bird escapees, tracing their origin back to the Mongolian emperor Kublai Khan’s hunting expedition (ca 1290 A.D.), which included 10,000 falconers carrying a “vast number of gyrfalcons, peregrine falcons and sakers” (*7, 11*). Based on criteria that include head and plumage patterns, Altai region falcons are informally classified into saker-like and gyrfalcon-like falcons (*12*), which is the neutral definition we follow in this study, as it does not presume their taxonomic status. Nonetheless, without a robust and comprehensive classification of Altai falcons *(F. altaicus)*, conservation efforts would be impeded, misguided and sometimes detrimental (*13*–*16*). Being conflated with the saker falcon, whose range extends from Western Asia to Eastern Europe, the Altai region falcons may lack the due conservation attention and informed management it requires.

Previous studies, based on conventional genetic markers (microsatellites and partial mitochondrial sequences) have not conclusively resolved the phylogenetic relationship between the saker and the gyrfalcon (*17*). While the sequenced genome of the saker falcon has allowed a more detailed insight into its genetic and the demographic history, it was not without limitations. Similar to the peregrine falcon, the saker’s genome size was found to be approximately 1.2 Gbp in length, encoding about 16,200 genes (*18*). However, these previous genome assemblies, along with the recently sequenced prairie falcon (*19*), were generated using short-read sequencing technology, resulting in highly-fragmented contigs that are prone to misassemblies and fall short of capturing whole chromosomes and the other high-quality standards proposed by the Vertebrate Genomes Project (VGP) (*20*). The resolution of the genomes of non-model organisms that have relatively high heterozygosity between haplotypes has been a challenge for short-read sequencing technologies and assembly algorithms, contributing to high fragmentation and/or misassemblies (*21, 22*). The resultant assemblies are highly fragmented, have lots of gaps and poor resolution of structural variations, which can have critical conservation, adaptive, and phylogenetic implications (*23*–*25*). As a result, the annotated dataset may contain missing, partial and/or misassembled genes, coding sequences (CDSs) and gene boundaries (*26*–*29*). Capturing such inaccessible genomic information by using long-read sequencing and other complementary technologies would allow for more accurate comparative studies on population, into hybridisation and speciation (*25*), and inform the conservation and management strategies. Here we generated a high-quality, chromosome-level VGP assembly of the gyrfalcon, followed by a comprehensive population analysis of gyrfalcons, sakers, Altai region falcons, peregrines, lanner and their hybrids, and used it to address the taxonomic status of the puzzling “Altai falcon” (*F. altaicus*).

## Results

### High-quality chromosome-level annotated gyrfalcon assembly

Short-read based genome assembly projects often sequence homogametic individuals (i.e. males in birds and females in mammals) to avoid the issues that arise from attempting to assemble the highly repetitive heterogametic W or Y chromosomes (*30*–*33*). To represent both W and Z chromosomes we chose a female gyrfalcon instead (females are the heterogametic sex in birds). We generated 62 Gbp of Pacific Biosciences continuous long reads (CLR; 51.6X-coverage) and 121 Gbp of Illumina short-read raw data (101.3X), 150.87 Gbp of linked reads using 10x Genomics (125.7X), 120.83 Gbp of Arima Hi-C data (100.7X) and 345 Gbp of Bionano Optical map data (287.5X). The data were assembled following the VGP pipeline 1.6 (*20*). Briefly, contigs were generated with FALCON unzip, which were then scaffolded and polished with 10x Genomics linked reads, followed by further scaffolding with Bionano optical genome maps and Hi-C linked reads to chromosome-level. This resulted in a contig N50 of 15.8 Mbp and NG50 of 60.5 Mbp, and a scaffold N50 and NG50 of 91.1 Mbp of the primary haplotype (Table 1). The estimated K-mer-based Quality Value (QV) of the final reference assembly is 40.6 with K-mer based completeness of 96.9%, and BUSCO completeness of 98.3% (2,543 Complete Universal Single-Copy Orthologs, with only 23 orthologs missing; Fig. S1). The number of gaps is 517. After manual curation guided by Hi-C maps, we identified 22 autosomes in addition to W (8,193,377 bp) and Z (84,945,525 bp) chromosomes, with a total genome size of 1,195,847,496 bp (Table S1). This reference assembly satisfies the VGP minimum metric values (*20*), and has a ∼500-fold increase in contiguity compared to the all previous short read genomes assemblies of any falcon species (Table 1). It was uploaded to NCBI and annotated with transcriptome data under accession number GCF_015220075.1.

**Table 1.** Summary statistics of the assembly parameters for the gyrfalcon reference genome generated in this study compared to the published draft genome assemblies of three falcon species (saker, peregrine and prairie). Abbreviations: C = contigs and S =scaffolds.

| Species | Common Name | Assembly Total size (Gbp) | Number of contigs (C), scaffolds (S) | N50 contig (C), scaffold (S) | Gaps within scaffolds | Ref. |
| --- | --- | --- | --- | --- | --- | --- |
| *Falco rusticolus* | **Gyrfalcon** | 1.20 | C: 746<br>S: 37 | C: 15.8 Mbp<br>S: 91.1 Mbp | 517 | This study |
| *Falco cherrug* | Saker | 1.18 | C: 56,956<br>S: 1,069 | C: 31.2 kbp,<br>S: 4.15 Mbp | 70,035 | Zhan et al. 2013 |
| *Falco peregrinus* | Peregrine | 1.17 | C: 61,119<br>S: 1,092 | C:28.6 kbp<br>S: 3.89 Mbp | 22,920 | Zhan et al. 2013 |
| *Falco mexicanus* | Prairie | 1.17 | C: not reported<br>S: 2,181 | S: 3.7 Mbp | 105,331 | Doyle et al. 2018 |

The *ab initio* annotation yielded 19,301 genes (including coding and non-coding), which are within the expected range of annotated genes in other published avian genomes, especially that of falcons (*18, 19*). Of these, 1,612 and 462 genes were on the Z and W chromosomes, respectively, including known sex-linked genes such as the *CHD1* (chromodomain-helicase-DNA binding) on both chromosomes, and Z-linked *DMRT1* (Doublesex and mab-3 related transcription factor 1). The assembled size (∼85 Mbp) and number of genes on the gyrfalcon’s Z chromosome are in line with the expected size and gene count of the highly conserved avian Z chromosome (*34, 35*). The final annotation of the assembly was done using the National Center for Biotechnology Information (NCBI) Eukaryotic Genome Annotation Pipeline. The analysis identified 15,894 protein-coding genes, of which 734 are Z-linked, and 186 are W-linked (GCA_015220075.1). Although the number of genes is comparable to short-read based falcon assemblies, the high contiguity and base quality of the gyrfalcon reference genome presented here, coupled with the significant reduction in gaps, yielded more complete genic and intergenic sequences, including phased euchromatic sequences. For example, the NCBI Annotation Release 102 for the short read saker falcon assembly (accession: GCF_000337975.1) identified 15,025 protein-coding genes, out of which 1,319 partial coding sequences (CDS). In contrast, NCBI annotation of the gyrfalcon annotated 41,414 CDSs, of which only 105 are partial CDSs.

### Admixture in wild falcon populations in Mongolia and the identification of a highly differentiated W chromosome

Population structure and admixture analyses were carried out using three whole genome sequence datasets of high-quality autosomal, Z- and W-linked Single nucleotide polymorphisms (SNP) (Table S2). Principal components analysis (PCA) using 191,122 autosomal SNPs separated the sakers (SF) and gyrfalcons (GF) along the first axis (Fig. 1A). Both falcons from the Altai region, the saker-like (SLF) and gyrfalcon-like (GLF), clustered closer to the species they morphologically resembled.

**Fig. 1.**
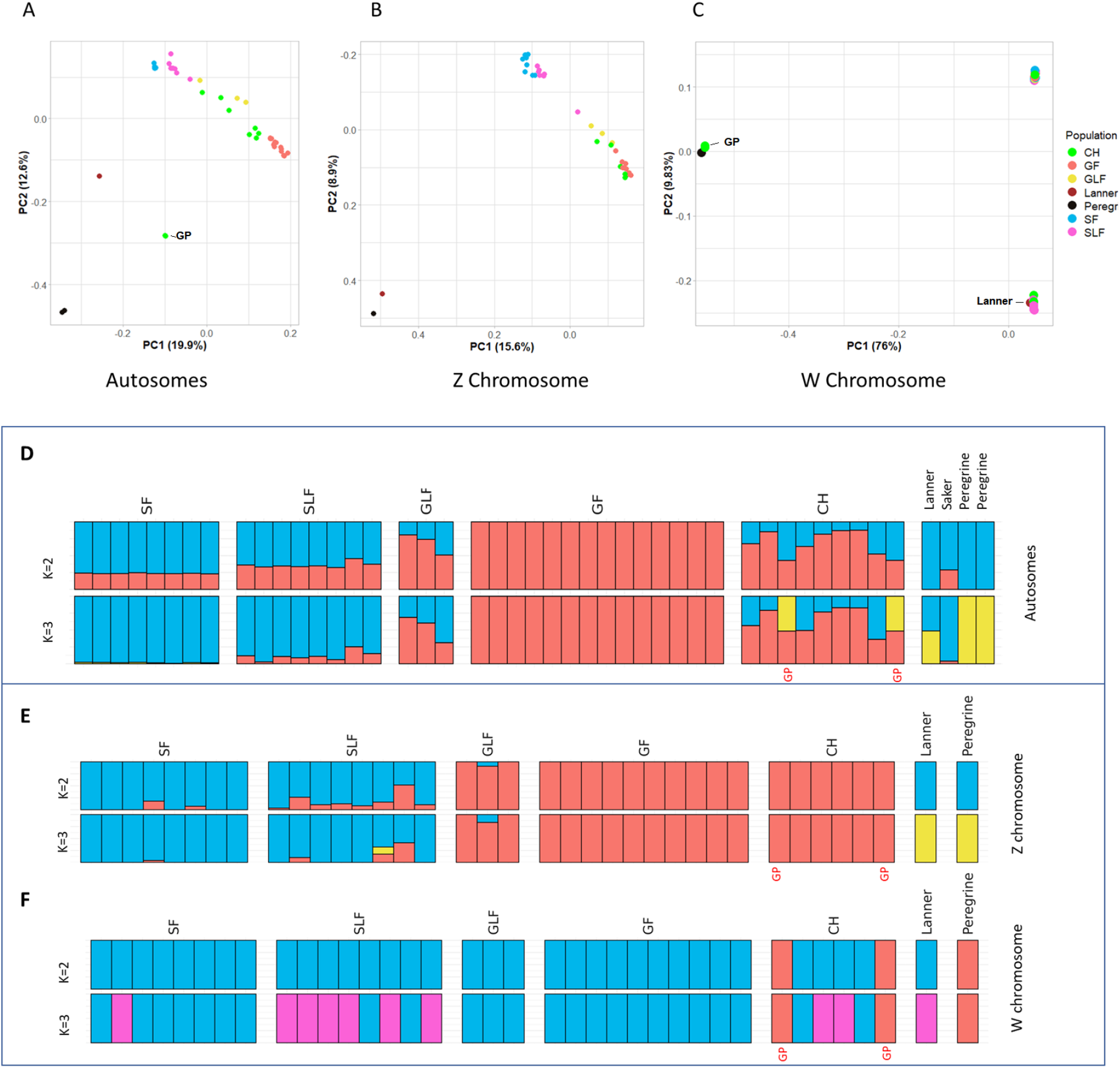
Population structure analysis of 46 falcon genomes. (**A)**, (**B)** and (**C)** shows Principal Component Analysis (PCA), based on autosomal, Z and W-chromosomal SNP datasets, respectively. Individual falcons are represented by a circle and color coded by population. **(D), (E)** and **(F)** are ADMIXTURE analyses, where each bar represents the ancestry of an individual falcon using autosomal, Z- and W-chromosomal SNP datasets, respectively, at a specific population number (K value). A single color denotes a pure lineage, while admixed individuals show multiple colors based on their ancestral groupings. The populations are North American gyrfalcons (GF) and sakers (SF). Falcons from the Altai region are designated gyr-like falcons (GLF) or saker-like falcons (SLF). Commercial Hybrids (CH) including gyrfalcon-saker and gyrfalcon-peregrine (labeled with GP). The publicly available genomes of one saker and two peregrines were added to the analysis. See Table 2 for details on individual samples.

For the sex chromosomes, the Z-linked dataset based on 4,975 SNPs (Table S2) reflected a population differentiation similar to the autosomal dataset, with a few notable exceptions (Fig. 1B). The Z chromosome of both the lanner and the peregrine appears to be highly similar to each other, while dissimilar to both the gyrfalcon’s and the saker’s Z chromosome (Fig. 1B). The most striking result was the distinct clustering on the W chromosomes (Fig. 1C). Only females were included in this analysis, since males do not have a W. The W-chromosome PCA (with 60,441 high-quality SNPs) revealed three demonstrably distinguished clusters: 1) Saker falcon (SF), gyrfalcon (GF) and gyr-like Altai falcon (GLF); 2) The majority of saker-like Altai falcons (SLF), along with the lanner, and 3) Peregrines and gyrfalcon-peregrine hybrids (Fig. 1C, labeled “GP”).

The PCA results were corroborated by the ancestral composition results (ADMIXTURE) (Fig. 1, D to F). The autosomal dataset was best represented at K=2, i.e. 2-population level, with a cross-validation (CV) value=0.54, and clearly separated gyrfalcons (GF) and sakers (SF) into two groups. All GF individuals showed a maximal gyrfalcon ancestry fraction (Q_G_=0.99999), while all SF individuals showed zero gyrfalcon ancestry fraction (Q_G_=0.00001) (Table S3). Based on these findings, we consider both of these populations as representatives of their corresponding species. In contrast, at K=2, SLF revealed notable levels of admixture between saker and gyrfalcon (Q_G_ ∼ 15%). The three GLF individuals also showed significantly higher gyrfalcon ancestry levels, up to 94% Q_G_, admixed with saker (Fig. 1D). The lanner appears to share genetic resemblance with both the sakers and the peregrines, but not with the gyrfalcons.

The ADMIXTURE structure of both sex chromosomes were also found to be concordant with their corresponding PCA analyses. The population structure based on the Z-chromosome generally resembles the autosomal patterns (Fig. 1, A, B, D, and E). The W-chromosome’s ADMIXTURE patterns at K=2 separate the peregrines from the rest of the falcons (Fig. 1F). At K=3, the SLF population seems to predominantly carry a distinct W-chromosome haplotype, which is also carried by the lanner. This “Altai haplotype” is different from the saker’s and the gyrfalcon’s. At K=3, two individuals, both gyrfalcon-peregrine hybrids (Fig.1F, labeled “GP”), form a distinct haplogroup along with the peregrine, as they carry a peregrine’s W chromosome from the mother side (Fig. 1F), and the gyrfalcon’s Z chromosome from their father’s side (Fig. 1E). In addition, our analysis of the Z chromosome of female hybrids revealed that gyrfalcon-saker hybrids tend to have a gyrfalcon sire (Fig. 1E, labeled “CH”). This confirmed the reported common practice among captive-falcon breeders where they use gyrfalcons as sires to increase the phenotypic resemblance (i.e. commercially valuable traits such as size and build) of the hybrid offspring to the gyrfalcons. This phenomenon is referred to as “paternal effect”, as falcon sires contribute more sex-linked alleles to the offspring than the falcon dams (*36*).

In order to examine the nature of the admixture of SLF and GLF and discern incomplete lineage sorting (ILS) from hybridisation, we performed a D-statistic test (or ABBA-BABA test, Fig. 2). The D-statistic test is a parsimony-like method for detecting gene flow, differentiating it from Incomplete Lineage Sorting (ILS) (*37*). The D-statistic classifies alleles as ancestral (‘A’) and derived (‘B’) across the genomes of four populations; two ancestral, one “admixed” and one outgroup. In this case, the two ancestral populations are the gyrfalcons and the sakers, the potentially admixed populations (SLF or GLF) and the outgroup is the peregrine. In the case of ILS alone, the number of discordant alleles (SNPs) grouping the ancestral populations with admixed populations should be roughly the same, i.e. BABA ≃ ABBA. However, a significant difference (|z| score > 3) between ABBA and BABA indicates that one of the ancestral populations shares more derived alleles with the potentially admixed population than expected by chance, implying introgression and gene flow between these two populations (*38*). Using autosomal and Z-linked SNPs, both populations from the Altai region show significant gyrfalcon introgression. However, GLF shares more derived alleles with the pure gyrfalcons compared to SLF (Fig. 2). GLF also appears to have a W chromosome that is closely related to pure gyrfalcons (Fig. 2B). On the other hand, the majority of SLF seem to have a distinct W chromosome, as evident by the negative D (Z score = −9.7) (Fig. 2A), which indicates that the W-chromosome of sakers and gyrfalcons share more derived alleles with each other than either one of them does with the Altai W-haplotype.

**Fig. 2.**
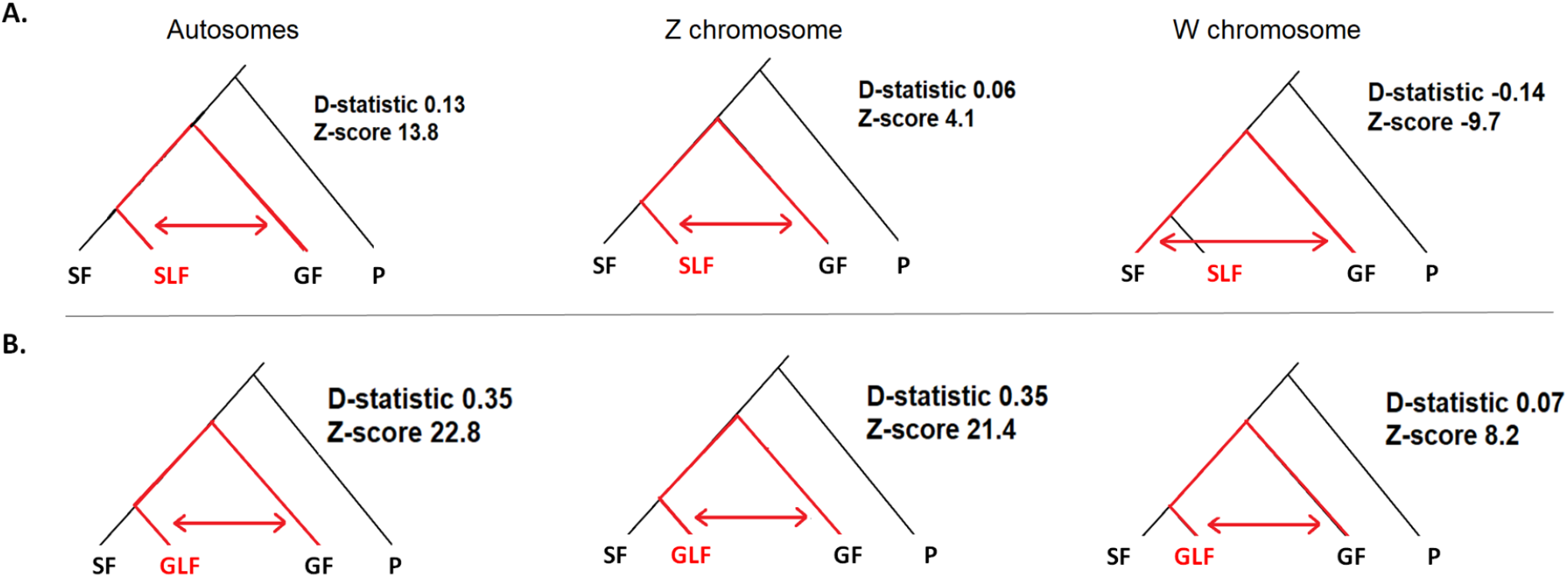
D-statistics analysis for (**A**) Altai saker-like falcon, and (**B**) Altai gyr-like falcon estimated using three datasets; autosomes, Z-chromosome, and W-chromosome. The significance level is calculated using Z score. The populations are saker (SF), North American gyrfalcon (GF), Altai gyrfalcon-like (GLF), Altai saker-like (SLF) and Peregrines (P) as an outgroup. The red arrow represents the excess allele sharing.

### Altai haplotype split 0.422 MYA from saker and gyrfalcon haplotypes

Next, in order to to estimate the divergence time of the Altai haplotype from the other closely-related falcon species, we assembled complete mitogenomes *de novo* for 35 individuals and generated a time-calibrated phylogenetic tree (chronogram) (Fig. 3A). We also included additional falcon reference mitogenomes retrieved from NCBI (Accession numbers: common kestrel NC_011307.1, American kestrel NC_008547.1, merlin KM264304.1, saker NC_026715.1, peregrine 1 NC_000878.1, peregrine 2 JQ282801.1 and gyrfalcon KT989235.1). We constructed a UPGMA phylogenetic tree using an alignment of W-specific variants (4257 SNPs) in 36 female falcons (Fig. 3B). The analyses of the mitochondrial phylogeny corroborated the distinct W haplotype observed in saker-like Altai falcons (Fig. 3, A and B), consistent with both being maternally-inherited in birds. Peregrines diverged from hierofalcons around 2.77 MYA, a result consistent with previous estimates (*39, 40*). Hierofalcon is a complex of species that includes the saker (*F. cherrug*), gyrfalcon (*F. rusticolus*), lanner (*F. biarmicus*), luggar (*F. jugger*) and the Australian black falcon (*F. subniger*) (*17*). The hierofalcon group was further resolved into two main clades: one comprising the majority of SLF individuals along with the lanner falcon, and another comprising SF, GF and GLF. The two clades separated ∼0.422 MYA. Two of the 8 SLF individuals (SP42 and SP35) clustered with the sakers (SF), whereas SP32 clustered as an SLF. The divergence time between GF and GLF, on one hand, and SF on the other was much more recent and estimated to be around 0.109 MYA. Our results also show that the gyr-like Altai falcons carry a haplotype that split 0.033 MYA from the North American gyrfalcon haplotype (Fig. 3A).

**Fig. 3.**
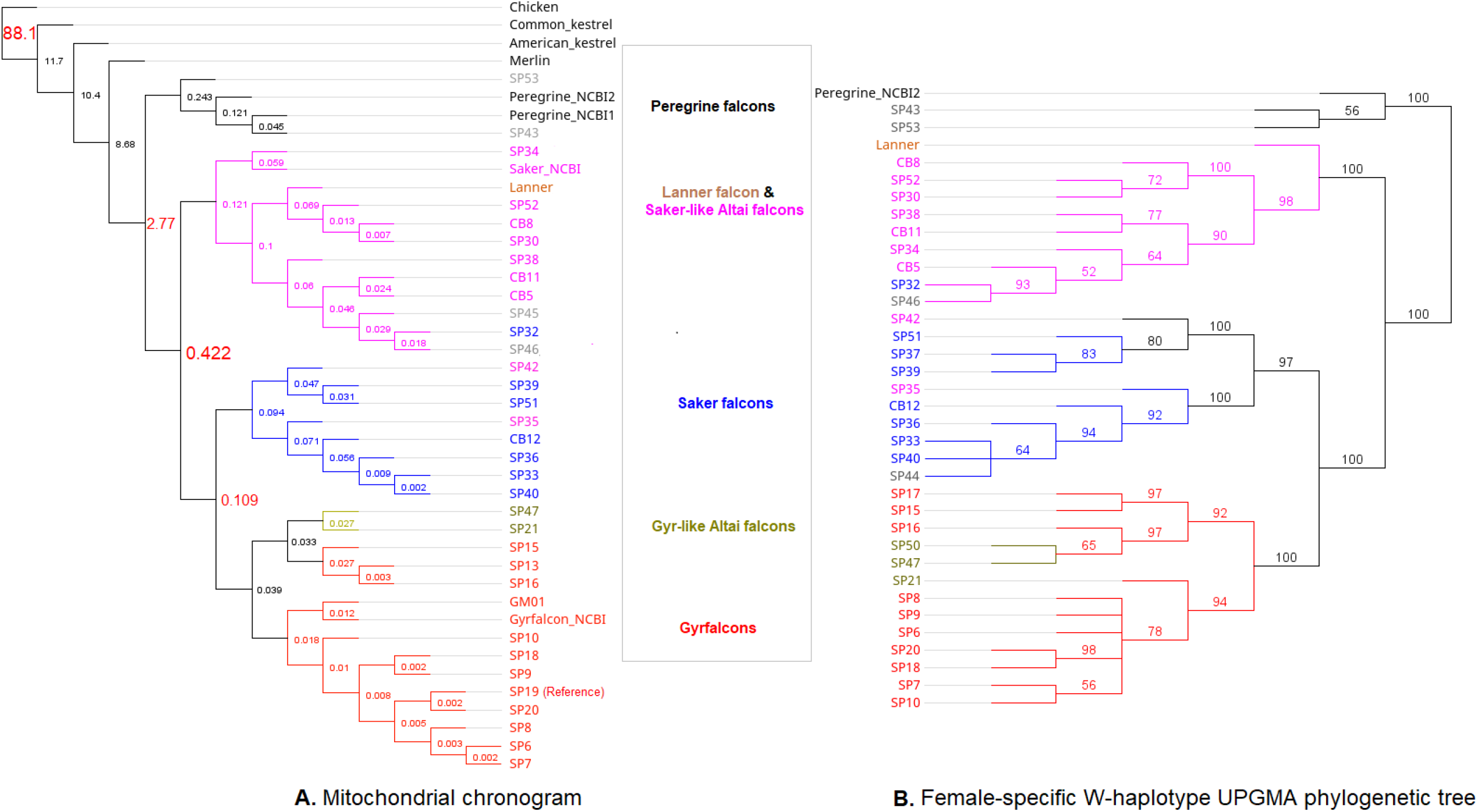
Divergence times of multiple falcon species and populations. (**A)** Chronogram (time-calibrated phylogenetic tree) of 42 falcons based on whole mitogenome (mtDNA) sequences (excluding the control region, CR) with chicken mtDNA as an outgroup, generated using Beast2. The sequences were aligned using CLUSTALW and nodes labeled with divergence time in million years ago (MYA), based on the estimated divergence time between chicken and falcons of 88 ± 10 MYA. (**B)** UPGMA phylogenetic tree using an alignment of W-specific variants (4257 SNPs) in 38 female falcons. The tree was constructed using HKY genetic distance model with a bootstrap value of 100, which is shown as a branch label. The red cluster denotes gyrfalcons (GF) and the yellow denotes gyr-like Altai falcons (GLF), the blue cluster represents sakers (SF), and the purple cluster represents saker-like Altai falcons (SLF). SP19 mitogenome is the reference mitogenome that was assembled in this project. Gray labels refer to hybrid individuals that were added to validate the Y chromosome haplotype results. sequences (labeled with the species name) were retrieved from NCBI (Accession numbers: common kestrel NC_011307.1, American kestrel NC_008547.1, merlin KM264304.1, saker NC_026715.1, peregrine 1 NC_000878.1, peregrine 2 JQ282801.1 and gyrfalcon KT989235.1). A full tree with confidence intervals of the age estimates can be found in Fig. S2.

To uncover the dynamics of the demographic history of falcon populations, we performed pairwise sequentially Markovian coalescent (PSMC) utilizing the heterozygous autosomal sites identified in their genomes based on their autosomal data (Fig. 4). Our analysis showed inferred fluctuations in the effective population sizes (Ne) of four species of falcons along with saker-like Altai falcon from 5 MYA to 10,000 years ago. We also identified a proto-hierofalcon as the most common recent ancestor of lanners, gyrfalcons and sakers, which appear to diverge from the ancestral peregrine falcons around 2-3 MYA (Fig. 4). This finding is consistent with the divergence time inferred using mitogenomes (i.e. 2.77 MYA, Fig. 3A). More specifically, the population of the proto-hierofalcon seems to expand until around 1 MYA, when the lanner splits from the rest of the hierofalcon group, and continued to expand well into the Last Glacial Period (LGP, 115,000 - 11,700 years ago; Fig. 4) when the global climate was colder. Meanwhile, the ancestral population of the gyrfalcons, sakers and Altai region falcons (SLF and GLF) appears to have experienced a bottleneck that lasted until around 200,000 years ago. Afterwhich, the saker population, like the lanner, demonstrates a drastic expansion, overlapping with LGP (Fig. 4). On the other hand, the gyrfalcon (both North American and GLF) appears to have never recovered from the bottleneck, maintaining a low, but steady effective population size until 10,000 years ago, even during the LGP. This may reflect an early adaptation of the gyrfalcons to cold climates, which also may have restricted the expansion of its population. Gyrfalcons are considered specialist predators in terms of diet and habitat. Unlike other falcons, gyrfalcons are a resident species in the Arctic tundra, adapted to the steeply cold weather, and specialized in preying on rock ptarmigan (*41, 42*). These factors may have contributed to the stabilization and limitation of the gyrfalcon population throughout its demographic history. The peregrine population appears to have experienced a bottleneck after they diverged from the proto-hierofalcon, around 2 MYA, while maintaining a steady effective population size, until around 500,000, when the peregrine population seems to have expanded, before declining around 50,000 years ago (Fig. 4).

**Fig. 4.**
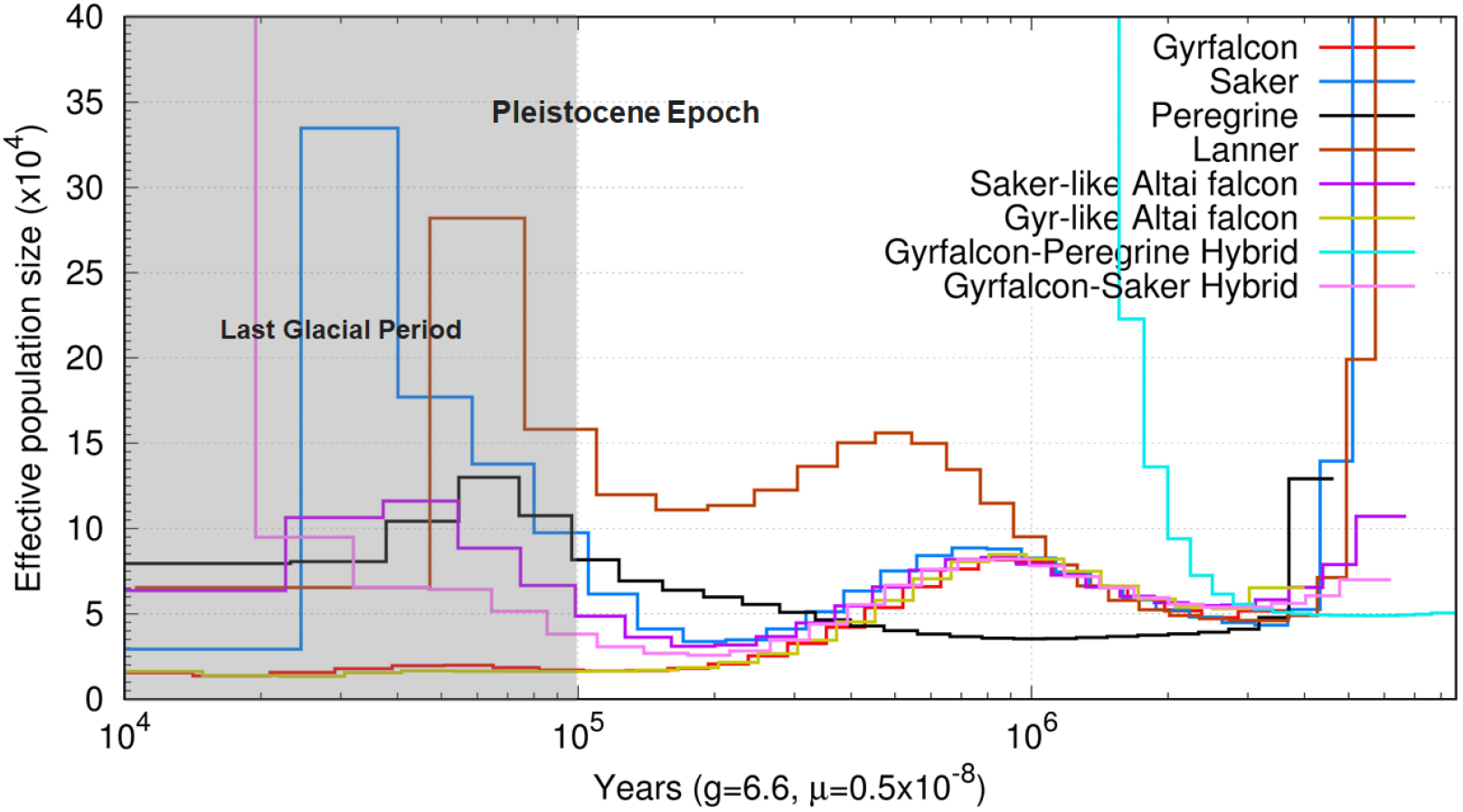
Pairwise sequential Markovian coalescent (PSMC) analysis of the autosomal data based on the heterozygous sites, showing the demographic history of four falcon species and two F1 hybrids and one Altai saker-like falcon. It demonstrates the fluctuation of effective population size from 6 million to 10,000 years ago. Time scale on the x-axis is calculated assuming a mutation rate of 0.5×10^−8^ per generation and generation time equal to 6.6.

It has been suggested that the spike observed in effective population size (Ne) of an F1 hybrid represents the point at which gene flow ceased between two populations (*43, 44*). Using an F1 gyrfalcon-peregrine hybrid, the spike pattern indicates the maximum bound of divergence time between the most recent common ancestor of hierofalcons and ancestral peregrines. We estimated this spike between 2-3 MYA (Fig. 4), in support of the phylogenetic estimate previously reported (*18, 45*), as well as the mitochondrial-based divergence time estimated in this study, i.e. 2.77 MYA (Fig. 3).

### W chromosome genes influence speciation

The gyrfalcon’s W chromosome with a size of 25.6 Mbp is significantly larger compared to 9.1 Mbp in the chicken (VGP assembly, GCF_016699485.2) and larger than the 20 Mbp W chromosome of the Zebra finch (VGP assembly, GCF_003957595.2). The expanded size of the falcon’s W was also reflected in its relatively high gene content with 186 protein-coding genes (Table S4), compared with 90 found on the chicken’s W chromosome and 165 protein-coding genes found on the Zebra finch’s W. Functional analysis shows that gyrfalcon’s W-linked genes are involved in multiple reproductive and development pathways including the Gonadotropin-releasing hormone receptor pathway, angiogenesis, fibroblast growth factor (FGF) and transforming growth factor β signaling pathways (Fig. 5A and Table S5). A few of these genes appear to be exclusive to the gyrfalcon’s W chromosome; *NEK5, ERVV-2, WASHC3, ERVW-1, ATP7B, VPS36, ALG11, SEC11C, RX2, ERVFRD-1* (2 copies) (Table S4).

**Fig. 5.**
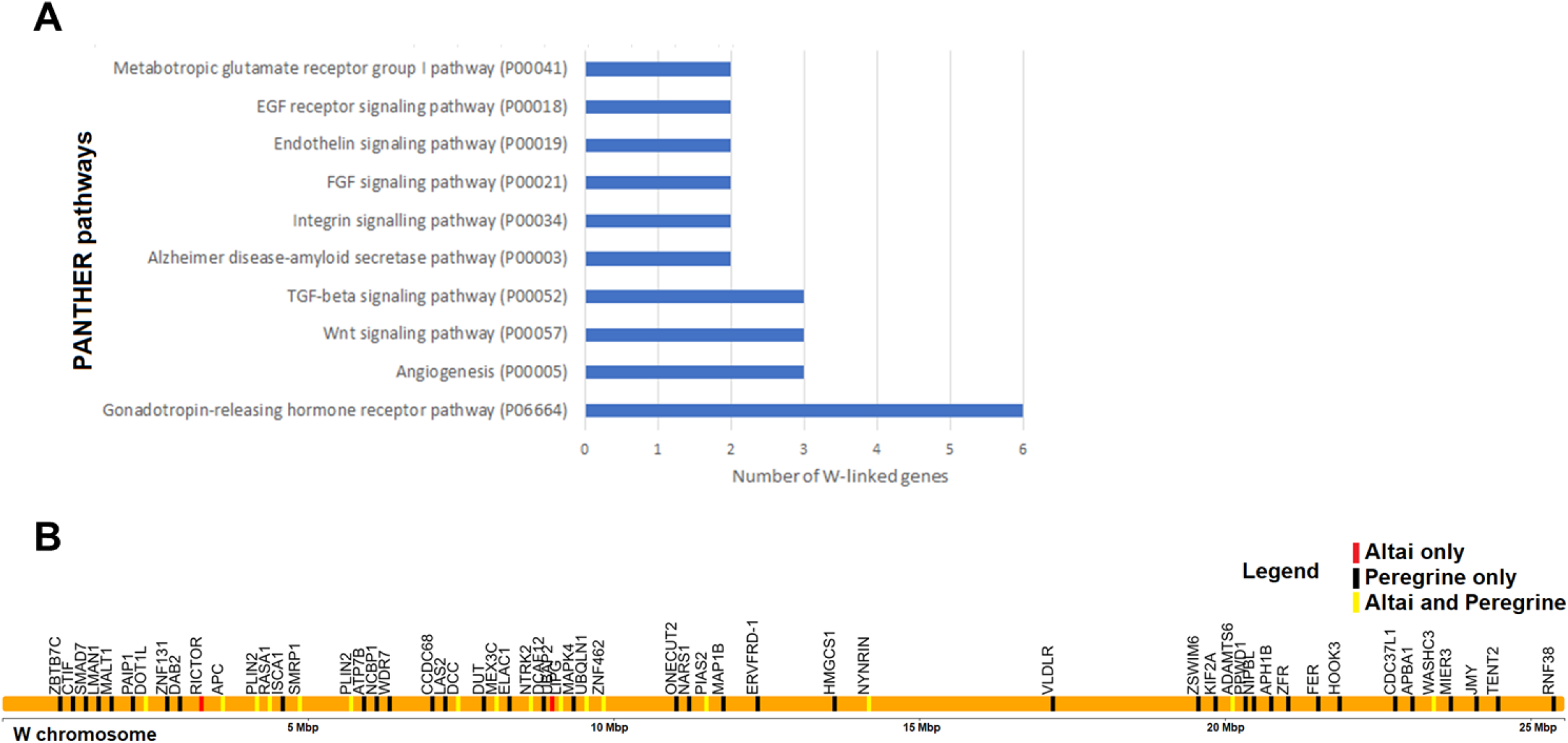
Functional Analysis of the W chromosome in falcons. **(A)** Number of W-linked genes involved in critical reproductive, physiological and pathological pathways using The PANTHER (Protein ANalysis THrough Evolutionary Relationships) Classification System. The top 10 PANTHER pathways are shown here. For a complete list see Supplementary Table S4. **(B)** W-linked genes affected by moderate or high impact mutations exclusive to peregrine, Altai, or found in both.

Some of these genes are also missing from the chicken and the zebra finch genomes, such as *ERVFRD-1*, which encodes syncytin-2, a protein associated with trophoblast development in humans (*46*). Many other W-linked genes belong to the endogenous retroviruses (ERVs) family including *ERVFRD-1, ERVW-1, NYNRIN, ERVK-5, ERVV2*. ERVs family of genes are thought to play a role in reproduction, female-biased mutational load and reproductive isolation (*30, 47*). Moreover, ERVs were found to function as species-specific enhancers in the germline gene expression, influencing spermatogenesis and possibly contributing to speciation (*48*). *NTRK2*, a gene associated with body weight in humans and mice (*49*), was found to have two copies on the gyrfalcon’s W chromosome. While it has gametolog on the Z chromosome, the two W-linked copies may suggest a possible overexpression in females that may contribute to the reverse sexual dimorphism (RSD) known in falcons, where females are larger than males. Another W-linked gene associated with body size is *NIPBL*, which, along with *NTRK2*, has been implicated in RSD in Chinese tongue sole (*Cynoglossus semilaevis*) (*50*). The presence of these genes on the gyrfalcons W chromosome may provide an explanation of the RSD in female falcons.

As the extent of the adaptive information provided by mitogenome variants is limited, we wanted to gain a deeper insight into the genetic basis underlying the selective variance within the W chromosome, and what may have led to the emergence of the Altai falcon haplotype. We looked at candidate genes under selection that may have contributed to the overall speciation trajectory of peregrines from the common ancestor with hierofalcons. Peregrines are a well-resolved species with a postzygotic reproductive barrier with hierofalcons. Out of 186 W-linked genes, 90 protein-coding genes contained a range of nonsynonymous variants in the peregrine W with predicted moderate (e.g. missense) to high (e.g. nonsense, stop-gain) impact (Fig. 5B and Table S6). These genes are associated with a range of phenotypes that could have contributed to the development of the peregrine as a separate species. For example, *SPINW*, a gene transcribed most prominently in ovarian granulosa and theca cells, that has been implicated in sexual differentiation in female chickens (*51*–*53*), appears to gain a stop codon in peregrines, affecting one of five transcripts of the gene (c.28C>T|p.Arg10*). The same mutation is also an intronic variant for the other transcripts. Another gene with a missense variant affecting both of its transcripts in peregrines (allele frequency 3/3) is *SMAD7* (c.148A>G|p.Ser50Gly). In female chickens, *SPINW* and *SMAD7* were reported to be upregulated in feather forming tissues (*54*), which is thought to contribute to their plumage sexual dimorphism. In contrast, sexual dimorphism in large falcons is restricted to size, where plumage is identical in both sexes, suggesting that large falcons may have lost the plumage-sexual dimorphism pathway, despite having genes involved in this pathway on the W chromosome. *THSD1*, an endocardial gene overexpressed in hearts in humans also has an alternative stop codon (c.1483T>C|p.Ter495Arg) in the peregrine. Other genes including *PLIN2W, UBAP2LW, DCAF1, NEK3, HINT1W, ARRDC3W, ISCA1* and *PIAS2* have also been identified with high impact variants (i.e. inactivating gene mutations) in the peregrine, along with the Altai-W haplotype in the case of the *LMAN1W* and *CHD1W* genes, that shifted the reading frame and introduced an alternative stop codons (Fig. 5B and Table S6). Notably, *HINT1W* is expressed in the developing gonads of female embryos at the critical time of sexual differentiation and has been suggested to play a role in avian sex determination (*55*).

Out of the peregrine’s W-linked genes under selection, the Altai W shares 25 of these genes with identical or similarly impactful variants as the peregrines (Table S6). This may hint at the ancestral W chromosome alleles of large falcons, given the split time of peregrines from Altai, compared to younger clades; i.e. saker and gyrfalcon. For example, both the Altai W-haplotype carriers (allele frequency 8/8) and peregrines (allele frequency 3/3) share a missense mutation (c.118C>T, p.Pro40Ser) in the only transcript of *ZNF462*, a gene associated with embryonic development and craniofacial and neurodevelopmental abnormalities in humans (*56*) and abnormal behavior in mice (*57*). A frameshift mutation (p.Gln771fs) identified in *CHD1W is* also shared between all the Altai-haplotype carriers and peregrine-W carriers with a predicted high impact (Table S6). Heterozygous missense variants in *CHD1* have been associated with developmental and facial structural changes (*58*). Speciation in falcons has also included gains in body size. Younger clades, including the largest and one of the youngest falcons, gyrfalcon, tend to be bigger and heavier compared to older clades such as the kestrels and the merlins. This has to do with the adaptation to new expansive ecologies and a wide range of prey, including larger ones such as the Willow Grouse (*Lagopus lagopus*) and Ptarmigan (*L. mutus*) (*59*). It is then expected that genes that are associated with body size, physiology, reproduction, and behavior to be under adaptive selection in a speciation trajectory (*60*). Indeed, *NTRK2, WASH3C, PLIN2W, PIAS2* and *MEX3C* genes are predicted to be impacted with high and/or moderate vartaints in peregrines and Altai W-haplotype carriers (Fig. 5B and Table S6). These genes are associated with vital structural, reproductive and physiological traits. For example, *WASHC3*, a gene associated with bill size in black-bellied seedcracker (*61*), shares a missense variant between Altai-haplotype carriers and peregrine-W carriers (Table S6). Another shared missense variant was found within one of *NTRK2* transcript*s*, a gene that is associated with the development of ovaries in mice (*62*) and litter size in sheep (*63*). Missense variants with predicted moderate impact were also identified in multiple ERVs, mainly *NYNRIN* and *NYNRIN-like*, which may be associated with the female reproductive physiology and potentially the development of a reproductive isolation (*30, 47*). Both copies of *PLIN2*, a gene associated with fat deposition in Peking Ducks (*64*), chicken (*65, 66*) and emu (*67*) are also impacted with a missense variant that is shared between Altai-haplotype carriers and peregrines. Moreover, the second *PLIN2* copy has a premature stop-gain variant (p.Leu437*) that is private to the peregrine W (Table S6). Perilipins, including *PLIN2*, also appear to play an important role in the physiological preparations for migration, including fat metabolism, in gray catbirds (*68*). Another gene, *MEX3C*, which is also linked to fat deposition and muscle development and whose function appears to be highly conserved in multiple species (*69*–*71*) is also predicted to be impacted in both peregrines and Altai-haplotype carriers (Table S6). *PIAS2* is reportedly associated with the immune response to viral infections in ducks (*72*). In *PIAS2* a missense variant was identified in Altai-haplotype carriers, and a nonsense mutation (stop-loss) was fixed in peregrines (Table S6). This gene being W-linked, and with variability among different species of falcons may contribute to a female-specific immunity. Significant sex-based variability in the immune system is also observed in many other species of birds (*73*–*75*). As apex predators, adaptive immunity plays an important role in the expansion into new niches, where potential prey may harbor novel pathogens. While the functional impact of these coding variants cannot be conclusively determined from genomic data alone, the pathways and traits with which they are associated make a strong case to consider them candidate variants in the speciation process. In general, the accumulation of mutations, under selective pressure, in genes linked to such critical traits could lead to incompatibilities with closely related populations, which would eventually create reproductive barriers, and ultimately the emergence of a new species (*60, 76*).

Our analysis also revealed that gyrfalcons and sakers share the same derived alleles in these loci. This demonstrates that the W chromosome is functionally conserved between these two species of falcons, implying a selective advantage.

## Discussion

Our results using the first high-quality chromosome-level genome of a falcon, help resolve the 200-year-old debate on the taxonomy of Altai falcons. As evident by the full assembly of the W chromosome and the mitochondrial genome, we have identified a distinct haplotype that seems to be predominant among Altai saker-like falcons. This Altai haplotype, along with the lanners, diverged more than 420,000 years from the common ancestor of saker and the gyrfalcon which, in turn, form a younger node that split around 100,000 years ago. The lanner falcon’s current distribution range is limited to Africa, Southern Europe, and fragmented and scattered points in the Middle East (*77*–*80*). Finding the lanner’s maternally-inherited haplotype in the Altai population of falcons in Mongolia, that share little physical resemblance with lanner, and with the lack of range contiguity to connect the two populations reinforces the idea of a complex demographic history of falcons. Moreover, we detected consistently similar patterns of autosomal admixture in Altai saker-like falcons, caused by a gene flow from both sakers and gyrfalcons. Both of these results suggest multiple hybridisation events. The first event would be a lanner-like population (as evident by the W haplotype), hybridizing with gyrfalcons. This would then be followed by a second event of consistent and repeated hybridization with sakers, which may explain the significantly higher saker-like ancestry, and saker-like Z chromosome detected in Altai saker-like falcons. If the scenario of the hybridisation events were in reverse order, the gyrfalcons would have donated their mitochondrial and W haplotype along with their autosomal markers, a conclusion that our results do not support in the case of the Altai saker-like population. However, this reverse order scenario seems to fit the admixture data of Altai gyr-like falcon individuals, as seen in the results. This alludes to the origin of gyr-like falcons as being indeed a gyrfalcon population that was introgressed by sakers, but maintained its gyrfalcon paternally- and maternally-inherited haplotypes, separating them from the Altai saker-like falcons.

East Asia, especially Mongolia, appears to be a hybridization hotspot for falcons, as our results reveal. If those admixture scenarios are true, it would be then expected, with a larger sample size, to detect admixed populations with maternal haplotypes of gyrfalcons, sakers and Altai region falcons. We would also expect to find autosomally admixed individuals with different ancestral proportions, reflecting both old and new hybridisations. Our results offer a glimpse of that, with one saker individual carrying an Altai maternal haplotype, and two Altai saker-like falcons carrying saker haplotypes. We also observed one gyr-like Altai falcon with a slightly higher saker ancestry (Fig. 1D, K=3), despite morphologically resembling gyrfalcons and carrying a gyrfalcon W and mitochondrial haplotype (Fig. 1F and Fig. 3A). All things considered, however, these observations do not change the fact that there are at least three maternally distinct, yet interbreeding populations of falcons in Mongolia; the gyrfalcons, the sakers, and the distinct Altai-haplotype falcons. As our data has shown, morphological appearance alone does not provide conclusive evidence of the taxonomy of these closely related falcon populations. This further highlights the importance of considering the autosomal admixture patterns along with the maternally-inherited haplotypes in discerning the taxonomy of Altai region falcons. While the designation of the gyrfalcons (including GLF) and sakers species is in line with the maternally-inherited haplotypes they carry, the Altai W-haplotype carrying falcons should also be designated as “Altai falcons” accordingly.

Proving homoploid hybrid speciation in animals is challenging, but with the aid of whole genome data, cases in birds and fish have been reported, where a hybrid lineage arises from interbreeding of two divergent species (*81, 82*). In the case of the genomically mosaic Altai falcons, the W chromosome appears to also play an important role in speciation, especially due to its maternally-inherited nature. By looking at the divergence of the peregrine W chromosome from the gyrfalcon’s and saker’s, we gained an insight into some of the genes that underwent selection, and hence the physiological, morphological and reproductive processes that contributed to the peregrine’s speciation trajectory. We then compared these W-linked genes to the Altai W chromosome to assess how functionally divergent it is from the older peregrine W haplotype and younger gyrfalcon-saker W haplotype. This potential could only be utilized by examining a fully-assembled W chromosome, instead of relying solely on mitochondrial analysis. Our results showed that not only does the Altai W chromosome share some of the functional mutations with the peregrine, but also that the impacted genes appear to be associated with vital pathways that could play a potential role in speciation. Considering the shared W-linked variants between the Altai and the peregrine, based on the D-statistics, phylogenomics and the closer examination of the shared functional variants, we find that Altai W is an older lineage and resembles the ancestral hierofalcon’s W haplotype. On the other hand, the saker and gyrfalcon, which split later, and share a functionally similar W haplotype, are shown to accumulate advantageous derived alleles in multiple W-linked genes. This demonstrates the impact of the female-biased gene flow on the trajectory of Altai falcon speciation process.

Our results suggest that the currently accepted falcon taxonomic classifications, mainly conflating the Altai falcons with sakers, should be revisited. If both gyrfalcons and sakers maintain their status as two distinct species and conservation units, despite sharing the highly similar sex-chromosome haplotypes and producing fertile offsprings (*6, 36*), then a fortiori the taxonomic status of Altai falcons should be reconsidered as well. The Altai falcons appear to be mosaic of multiple ancestries; they are predominantly of autosomally admixed origin, with a generally consistent saker and gyrfalcon ancestral proportions. They also carry a W haplotype that clusters with the lanner falcon and is separate from the ancestor of the gyrfalcon and the saker. Any autosomal traces of the lanner-like ancestry seems to be lost to the multiple subsequent admixture events with sakers and the gyrfalcons. Moreover, not only does the Altai W-haplotype split 0.42 MYA, but also it contains coding variants with implications on reproductive and structural traits. These multiple lines of evidence, both genetic and phylogenetic, support the upgrading of the Altai falcons (*F. altaicus*) to a species level.

Since the suggested species status of Altai falcons is based on phylogenetic and functional bases, it offers an essential foundation to inform conservation and management efforts. The focus of falcon conservation resources and projects currently ongoing in Mongolia, where Altai falcons reside, may need to be re-evaluated based on the current population trends of the Altai falcon, which is currently conflated with the saker falcon. The sakers, whose range overlaps with Altai falcons and extends as far as western Europe, may warrant their own separate conservation programs. In 2017, two populations of Sumatran orangutans (*Pongo abelii*) were recognized as two distinct species following extensive analyses (*83*). The new species, *P. tapanuliensis*, was found to have fewer than 800 individuals, calling for further research that focused on the newly identified species as a separate unit of conservation. A recent analysis revealed that the newly described orangutan species is perhaps the most threatened great ape in the world (*84*). This conclusion, and the subsequent recommendation for management and conservation, would not have been possible for this new species had it still been conflated with the significantly more abundant, yet still endangered, Sumatran orangutan (*P. abelii*). We hope that our work will facilitate and enable a similarly focused attention towards Altai falcons (*F. altaicus*) as a distinct unit of conservation, separate from sakers and gyrfalcons. Such designation of Altai falcons as a separate species will help inform and guide the ongoing conservation efforts of falcons in Asia, and maintain the genetic identity of Altai falcons by preventing accidental introduction of Central Asian saker falcons into its ranges. One of the major implications of our results is the power of using high-quality reference genomes for population genomics, in addressing long-debated questions in biology, taxonomy, ecology and conservation. The success of approaches, including breed-and-release, is critically dependent on taxonomic classifications, hence the need for a robust approach to delineating phylogenies. Defining units of conservation is the cornerstone of any conservation endeavor, without which it could lead to unpredicted and sometimes harmful consequences. The current work demonstrates the importance of expanding the use of W chromosomes beyond phylogeny, to include fingerprints of divergence and its vast implications in non-model species for conservation purposes. With the current mass extinction of species underway, a comprehensive and robust approach should be followed to determine the taxonomic status of populations, which in turn helps prioritize the conservation resources (*85*).

## Materials and Methods Sampling and animal ethics

A total of 44 falcons were sequenced (Table 2). These included: 14 gyrfalcons from different populations and color morphs, eight sakers; 11 Altai falcons (originating from western Mongolia, and based on morphological traits designated as Altai-type falcons); one lanner falcon; and nine commercial hybrids (seven gyrfalcon-saker and two gyrfalcon-peregrine hybrids). The Canadian and Alaskan gyrfalcon individuals (n = 13) were captive-bred using pure-bred lineages, without intraspecific hybridisation, i.e. crossing with other gyrfalcon populations; one individual (GM01) was labeled North American as its intra-specific captive-bred lineage was not precisely determined (Table 2). All the gyrfalcons had previously been exported to Qatar for breeding in captive-breeding facilities where they later were sampled. The sakers and the lanner were sampled in Qatar, while the Altai falcons (n = 11) had been previously exported from Mongolia (classified as sakers following the accepted taxonomy) to Qatar, where they were sampled. The latter resembled the morphology of two distinct populations of “Altai-type falcons”; gyrfalcon-like and saker-like (*12*). The commercial captive-bred hybrid falcons and their ancestry and filial generation were acquired from the breeders. Blood samples (< 1 ml, each) from each bird were collected in 1.5 ml tubes containing potassium EDTA anticoagulant, at the Souq Waqif Falcon Hospital, Doha, Qatar. Sample collection was carried out under the official approval of the Ministry of Municipality and Environment in Qatar (Reference No. 2017/283748) and by trained veterinarians at the Falcon Hospital. The blood samples were only taken during routine diagnostics from falcons being checked for purposes unrelated to the research activity and the researcher had no influence on the timing or location of the medical check-up. These blood samples would have normally been discarded if the researcher had not asked for them. The Monash University Animal Ethics Committee (AEC) advised that such activity, considered as scavenging, does not require animal ethics approval.

**Table 2.** The falcons used in this study for sequencing. The table shows sample ID, species name, sex, plumage color as well as provenance. W refers to an individual of a wild origin, while CB refers to a captive-bred individual. (?) after the species name refers to taxonomically-ambiguous individuals, i.e. Altai-type falcons. The reference specimen is marked with *.

| Sample ID | Species name | Common Name | Sex | Color | Provenance |
| --- | --- | --- | --- | --- | --- |
| North American Gyrfalcons (GF) |  |  |  |  |  |
| SP19* | *F. rusticolus* | Gyrfalcon | F | White | Alaskan (CB) |
| SP6 | *F. rusticolus* | Gyrfalcon | F | Black | Canadian (CB) |
| SP7 | *F. rusticolus* | Gyrfalcon | F | Black | Canadian (CB) |
| SP8 | *F. rusticolus* | Gyrfalcon | F | Black | Canadian (CB) |
| SP9 | *F. rusticolus* | Gyrfalcon | F | Black | Canadian (CB) |
| SP10 | *F. rusticolus* | Gyrfalcon | F | Black | Canadian (CB) |
| SP13 | *F. rusticolus* | Gyrfalcon | **M** | White | Alaskan (CB) |
| SP14 | *F. rusticolus* | Gyrfalcon | **M** | White | Alaskan (CB) |
| SP15 | *F. rusticolus* | Gyrfalcon | F | White | Alaskan (CB) |
| SP16 | *F. rusticolus* | Gyrfalcon | F | White | Alaskan (CB) |
| SP17 | *F. rusticolus* | Gyrfalcon | F | White | Alaskan (CB) |
| SP18 | *F. rusticolus* | Gyrfalcon | F | White | Alaskan (CB) |
| SP20 | *F. rusticolus* | Gyrfalcon | F | White | Alaskan (CB) |
| GM01 | *F. rusticolus* | Gyrfalcon | **M** | White | North American (CB) |
| Saker falcon (SF) |  |  |  |  |  |
| CB12 | *F. cherrug* | Saker | F | Light brown | Qatar (W) |
| SP32 | *F. cherrug* | Saker | F | Brown | Qatar (W) |
| SP33 | *F. cherrug* | Saker | F | Brown | Qatar (W) |
| SP36 | *F. cherrug* | Saker | F | Brown | Qatar (W) |
| SP37 | *F. cherrug* | Saker | F | Brown | Qatar (W) |
| SP39 | *F. cherrug* | Saker | F | Brown | Qatar (W) |
| SP40 | *F. cherrug* | Saker | F | Brown | Qatar (W) |
| SP51 | *F. cherrug* | Saker | F | Dark Brown | Qatar (W) |
| Saker-like Falcons from the Altai Region (SLF) |  |  |  |  |  |
| CB8 | *F. cherrug* (?) | Saker-like | F | Light-brown | Mongolian (W) |
| CB11 | *F. cherrug* (?) | Saker-like | F | Silver | Mongolian (W) |
| SP30 | *F. cherrug* (?) | Saker-like | F | Black | Mongolian (W) |
| SP34 | *F. cherrug* (?) | Saker-like | F | Black | Mongolian (W) |
| SP35 | *F. cherrug* (?) | Saker-like | F | Brown | Mongolian (W) |
| SP38 | *F. cherrug* (?) | Saker-like | F | Grey | Mongolian (W) |
| SP42 | *F. cherrug* (?) | Saker-like | F | Grey | Mongolian (W) |
| SP52 | *F. cherrug* (?) | Saker-like | F | Black | Mongolian (W) |
| Gyr-like Falcons from the Altai Region (GLF) |  |  |  |  |  |
| SP50 | *F. cherrug* (?) | Gyrfalcon-like | F | Grey/Silver | Mongolian (W) |
| SP21 | *F. cherrug* (?) | Gyrfalcon-like | F | Brown | Mongolian (W) |
| SP47 | *F. cherrug* (?) | Gyrfalcon-like | F | Silver | Mongolian (W) |
| Commercial hybrids (CH) |  |  |  |  |  |
| SP11 | *Hybrid* | F1 Gyrfalcon-Saker | **M** | Black | Hybrid (CB) |
| SP12 | *Hybrid* | F4 Gyrfalcon-Saker | **M** | Black | Hybrid (CB) |
| SP44 | *Hybrid* | F4 Gyrfalcon-Saker | F | White | Hybrid (CB) |
| SP45 | *Hybrid* | F4 Gyrfalcon-Saker | **M** | Black | Hybrid (CB) |
| SP46 | *Hybrid* | F5 Gyrfalcon-Saker | F | Sliver | Hybrid (CB) |
| SP48 | *Hybrid* | F4 Gyrfalcon-Saker | F | white | Hybrid (CB) |
| SP49 | *Hybrid* | F3 Gyrfalcon-Saker | F | White | Hybrid (CB) |
| SP43 | *Hybrid* | F1 Gyrfalcon-Peregrine | F | White | Hybrid (CB) |
| SP53 | *Hybrid* | F1 Gyrfalcon-Peregrine | F | Black | Hybrid (CB) |
| CB5 | *F. cherrug* | Saker | F | Light brown | Unknown (CB) |
| Lanner falcon |  |  |  |  |  |
| Lanner | *F. biarmicus* | Lanner | F | Wildtype | Qatar |

### DNA Extraction, Library Preparation and Resequencing

High molecular weight genomic DNA (HMW gDNA) was extracted from all of the blood samples that were collected (Table 2) using Qiagen’s MagAttract Kit (Qiagen, Germany). Extracted DNA was quantified and its quality and integrity were assessed by using agarose gel electrophoresis and Qubit® 3.0 Fluorometer (Thermo Fisher Scientific, USA). For each sample, ∼1 µg of gDNA was used for preparing an indexed library for sequencing using the NEBNext® Ultra™ DNA Library Prep Kit from Illumina® (NEB, USA) following the manufacturer’s protocol. The gDNA was randomly fragmented to a size of 350 bp by mechanical shearing. Subsequently, the DNA fragments were end-polished, A-tailed, and ligated with the NEBNext adapter for Illumina sequencing, and further PCR enriched by P5 and indexed P7 oligos. The PCR products were purified (AMPure XP system) and the resulting libraries were analyzed for size distribution on an Agilent 2200 TapeStation System (Agilent, USA). DNA resequencing was carried out for all the samples (Table 2) at the Sidra Medicine, Doha, Qatar, using Illumina HiSeq X platform at an average coverage of 30X. Raw sequencing data were filtered and assessed for quality using FastQC 0.11.8 and Trimmomatic 0.39 (*86*). The lanner genome was sequenced using PacBio Sequel II system at a coverage of 25X at Vertebrate Genome Laboratory (VGL), The Rockefeller University, New York, USA.

### Reference Genome Sequencing

High coverage DNA sequencing was done of the SP19 individual. The reference sample was also sequenced using Illumina HiSeq 4000 (150 bp paired-end) at a 101.2x coverage at Novogene Genomics, Singapore. For long reads, PacBio Sequel System v5.0, using chemistry v2.1 was done at a 51.7x coverage; Bionano Optical Mapping (DLE-1 one enzyme non-nicking approach using the Bionano Saphyr instrument); and 10x Genomics libraries were sequenced, all at the Vertebrate Genome Laboratory (VGL), The Rockefeller University, New York, USA. Chromosome conformation capture (Hi-C) was performed at Arima Genomics, USA. All raw data were uploaded on VGP’s GenomeArk repository for raw data and assemblies (https://vgp.github.io/genomeark/Falco_rusticolus/).

### Reference Genome Assembly and Annotation

High quality reference genome assembly was generated using VGP methodology (*20*). For PacBio’s long reads, Canu 1.8 (*87*) and pb-assembly 0.0.6 (FALCON 1.3.0 and FALCON_unzip 1.2.0) (*88*) were used to generate a phased diploid and polished draft assembly. Thirty iterations of PacBio draft assemblies were generated to optimize the contiguity and phasing and decrease duplications, guided by the results of Benchmarking Universal Single-Copy Orthologs (BUSCO) analysis (Fig. S1). The 10x Genomics data was processed using the Longranger 2.2.2 software and subsequently employed to refine and scaffold the PacBio draft assembly using ARCS+LINKS Pipeline (*89, 90*), (Fig. 6). To further optimize the scaffolding and resolve misassemblies, Bionano optical mapping data were integrated using Bionano Solve v3.2.1. Hi-C data were used to further scaffold the assembly to chromosome level using the Arima Genomics mapping pipeline (https://github.com/ArimaGenomics/mapping_pipeline) followed by Salsa2 (*91*). Finally, to improve the base quality of the assembly, two polishing methods were utilized: First, using PacBio long reads, polishing was conducted using Arrow (included in SMRT Link v.7). In Arrow, a range of coverage cutoff values were investigated to optimize the polishing efficiency, and the best was found to be -x 5. Second, further polishing steps used 10x Genomics short linked-reads; two rounds of short read polishing were performed using Longranger align 2.2.2, FreeBayes (*92*), and BCFtools consensus (*93*), according to VGP 1.6 pipeline (*20*) (Fig. 6). To confirm the quality and structure of the chromosome-level assembly, a Hi-C map was generated using PretextView 0.1.2.

**Fig. 6.**
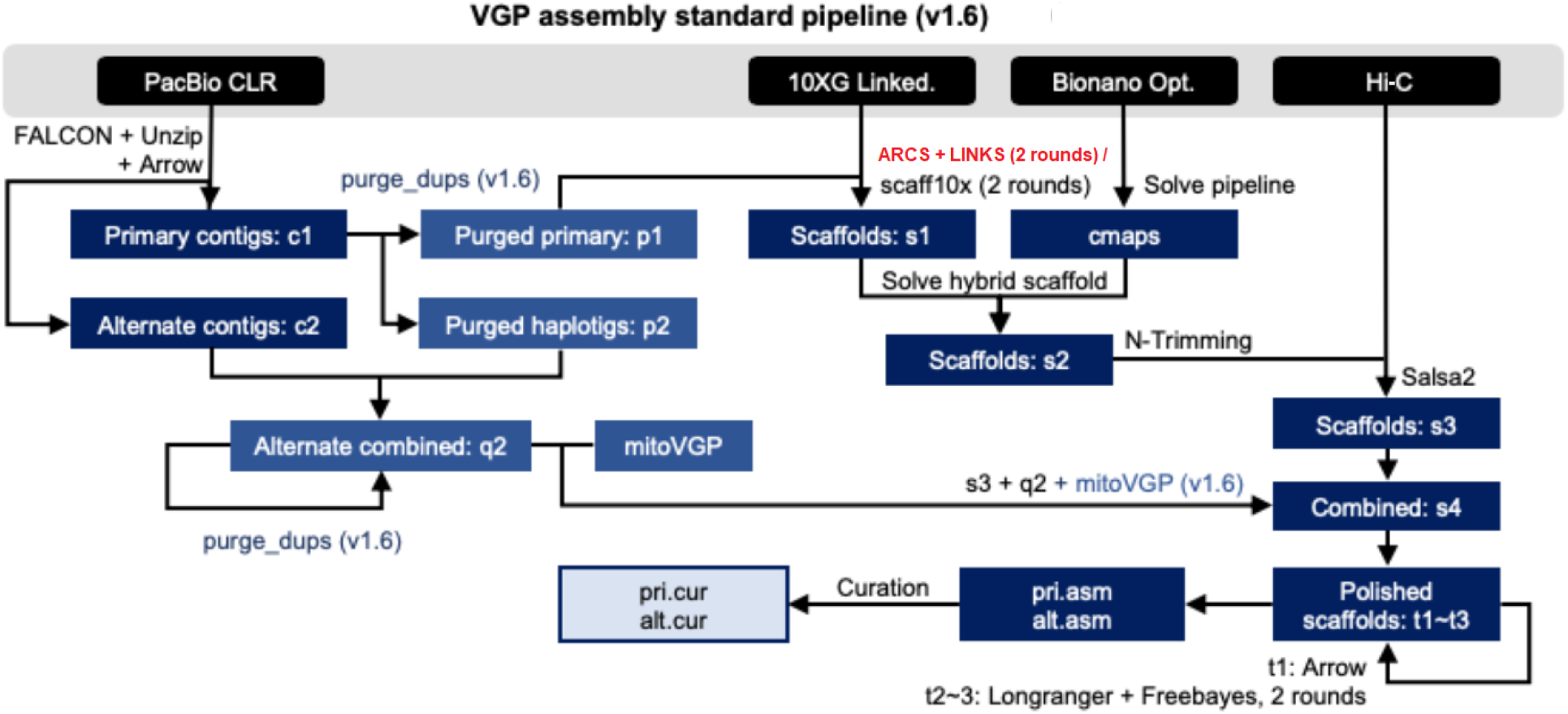
VGP standard assembly pipeline 1.6, with the modified step of ARCS+LINKS in parallel with scaff10x, used in this project. Abbreviations: c = contigs; p = purged false duplications from primary contigs; q = purged alternate contigs; s = scaffolds; t = polished scaffolds. Further details on the pipeline and instructions are available at https://github.com/VGP/vgp-assembly (*20*). In brief, the raw data from PacBio sequencing CLR technologies are assembled to contigs using FALCON + Unzip, followed by polishing using Arrow to increase the accuracy of the base calling. The primary contigs refer to the full genome, whereas the alternate refers to detected haplotypes within the diploid genome. The primary contigs undergo purging to remove artificial duplications within the assemblies. This is followed up by scaffolding using 10x Genomics linked-reads. The resultant scaffolded assembly is further upgraded to enhance the completeness and contiguity by integrating Bionano optical maps followed by Hi-C data to generate chromosome-level assembly. The resultant assembly is then polished, and gap-filled using the 10x Genomics data. The assembly that is generated was submitted for manual curation and further contiguity enhancement based on the Hi-C data.

All iterations were evaluated using quality assessment statistics including contig N50 and scaffold N50, and genome completeness using BUSCO v3 software (*94*). BUSCO provides quantitative measures for the assessment of genome assembly using a database of 2,586 near-universal single-copy vertebrate orthologs, included in the vertebrata_odb9 database. The Quality Value (QV) of each step of the assembly pipeline was estimated using Illumina short reads and 10x Genomics short reads using Merqury (*95*).

Draft annotation and gene prediction of the assembled genome was carried out using three *ab initio* pipelines: Augustus 3.3.2 (*96*), using chicken (*Gallus gallus*) as the model avian species, after identifying and masking repetitive DNA using RepeatMasker 4.0.9 (*97*). Functional annotation was done using Blast2GO (*98*), using integrated InterProScan and non-redundant protein sequences NCBI Blast (nr v5) databases. The transcriptomic database from NCBI, which included data from the *Falco* genus was used to validate the gene annotation as it is more comprehensive than other available approaches. A functional annotation of the gyrfalcon annotation using Gene Ontology (GO) enrichment analysis was performed to evaluate the completeness of the draft assembly and the annotation. The reference-quality, manually curated, and annotated assembly for the gyrfalcon is available at NCBI (GCF_015220135.1).

### Genetic variation, admixture and population structure

The paired-end short reads for the additional resequenced samples were mapped to the reference assembly of the gyrfalcon generated in this study (GCF_015220135.1) using BWA-MEM alignment software (*99*). Publicly available WGS data of a saker (PRJNA168071), two peregrines (PRJNA159791, PRJDB7811) and a prairie falcon (PRJNA436967) were retrieved from NCBI and added to the analyses for comparison and validation. The mapped reads were then sorted using SAMtools 1.9. SNPs were then filtered (Phred quality score ≥ 20 and a minimum depth of 10 reads) and called using FreeBayes, VCFtools and BCFtools. The generated VCF file was split into three representing autosomal-, W- and Z-linked datasets. For admixture and population structure analyses, the filtered SNPs were then pruned to remove the SNPs that were in linkage or close proximity using an r^2^ cut-off of 0.1. The QC process also involved filtering out SNPs with: a) very low minor allele frequencies (< 0.05); b) autosomal SNPs with frequencies that deviate significantly from Hardy-Weinberg Equilibrium (p-value <10-6 threshold); c) excess missing genotypes (SNPs with more than 99.9% of genotypes missing), and d) low confidence call score (<Q30) score. The QC filtration was done using PLINK software v.1.9 (*100*) using the following command lines:

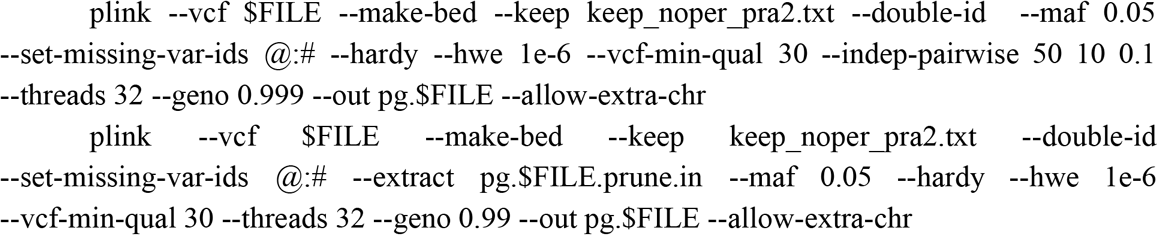

Using the PLINK software v.1.9 (*100*) a PCA was conducted to represent the variation in the populations and explore the population substructure across the autosomes and sex chromosomes. The same variation dataset was also used to estimate the likelihood of individual ancestries using ADMIXTURE 1.3 (*101*). The results were visualized using ggplot2 package in RStudio Cloud software (https://rstudio.cloud/).

To account for genetic drift and gene flow, TreeMix 1.13 was used, which infers patterns of population split using allele frequency (*102*). TreeMix was also used to calculate the F-statistics (F3 and F4) of all populations to estimate the mixing proportions of the admixture events (*103*). The D-statistic, also known as ABBA-BABA test, and its related f4-ratio ancestry proportion estimator were calculated for all populations using ADMIXTOOLS (*103*) through the *admixr* package version 0.7.1 (*104*) in RStudio (*105*).

### Mitochondrial genome assembly and phylogenetic analysis

A draft mitochondrial genome for the gyrfalcon was assembled using NOVOPlasty (*106*), by using the chicken mitogenome (NCBI accession No. KM433666.1) as a reference and a seed, and a k-value of 71. Whole-genome sequence datasets were used to assemble the full mitogenomes of 35 falcons from five populations using NOVOPlasty. Four mitogenomes (SP50, SP17, SP14, SP37) failed to assemble fully, possibly due to the relatively low sequencing coverage and computational requirements. The chicken (NCBI Accession KT626858.1) was used as an outgroup, and the analysis involved the inclusion of six falcon mitogenomes retrieved from NCBI: common kestrel (NC_011307.1), American kestrel (NC_008547.1), merlin (KM264304.1), saker (NC_026715.1), peregrine 1 (NC_000878.1), peregrine 2 (JQ282801.1) and gyrfalcon (KT989235.1). Following common practice, we excluded the control region (CR) due to its high variability, which could confound the phylogenetic signal (*107*–*109*). A neighbor-joining phylogenetic tree of the whole mitogenomes, excluding the control region, was constructed using Geneious Tree Builder following the Jukes-Cantor Genetic Distance Model and a consensus tree based on 1,000 bootstraps was generated.

### Demographic history

The falcon species divergence times were estimated on BEAST v2.6.2 (*110*) using StarBEAST2 module (*111*). We assumed a strict molecular clock, integrated analytical population size, and an HKY (four categories) nucleotide substitution model. For the priors, we selected the Calibrated Yule speciation model, and calibrated the molecular clock by assuming the age of divergence between Falconiformes and chicken 88 ± 10 MYA (*112*). Markov chain Monte Carlo chains (MCMC) were run for 60 million states, discarding the first 10% as burn-in. Convergence was inspected using Tracer v1.6, and a maximum clade credibility tree (chronogram) was built using TreeAnnotator v2.6.2 (*110*).

To reconstruct the population histories of the falcon species, we used the Pairwise Sequentially Markovian Coalescent (PSMC) model, which uses diploid sequences to infer piecewise-constant population size histories as a function of time (*113*). The raw sequencing reads were mapped to the reference, and heterozygous SNPs were called using SAMtools mpileup as in: samtools mpileup -C50 -uf ../ref.fa sample.bam | bcftools call -c | vcfutils.pl vcf2fq -d 10 -D 100 | gzip > sample.fq.gz. Autosomal SNPs were used to generate the PSMC files using the following command:

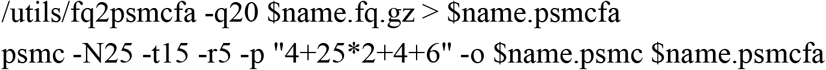

Finally, the PSMC files were plotted using the saker’s generation time of 6.6 years/generation according to previous estimates (*18*). A genomic mutation rate of 4.6×10^−9^ per base per generation that was obtained in a germline-based study for the collared flycatcher *Ficedula albicollis* (*114*) and has since been used as a reliable estimate for other birds as well (*115, 116*).

### Identification of candidate genes under selection and enrichment analysis

To identify candidate regions under selection, Hudson’s *F*ST (*117*) method was used. Hudson’s *F*ST was selected as it is independent of sample size, compared to other *F*ST calculation methods, including the classical Weir and Cockerham (*118*). *F*ST was computed using the script popgenWindows.py (github.com/simonhmartin/genomics_general) with a sliding window of 20,000 bp and a minimum of 100 genotyped sites. Only windows corresponding to the upper 0.5% of the empirical genome-wide distribution of *F*ST were considered as high-*F*ST outliers and labeled as candidate regions. The aim was to capture the gradual and subtle differences between the falcon populations, and highlight the highly differentiated genomic regions that could probably explain the adaptive phenotypes involved in their respective habitats and the rate at which they are diverging. SnpEff was used to annotate the variants within these genes, especially W-linked, and predict their impact (HIGH, MODERATE, LOW, and MODIFIER). Variants with predicted high or moderate impact were then investigated in depth and validated manually by examining the VCF file.

W-linked genes in the gyrfalcon were analyzed for functional roles using The PANTHER (Protein ANalysis THrough Evolutionary Relationships) Classification System version 17.0 (*119*)

## Acknowledgments

The authors would like to thank Al-Gannas Qatari Society, The Cultural Village Foundation-Katara, Doha, Qatar for their support and funding since the inception of the project. We also thank Souq Waqif Falcon Hospital, Qatar University, Sidra Medicine, Um Haish Falcon and Houbara Conservation Center, Genomics Facility of Monash University Malaysia and Vertebrate Genomes Laboratory, Rockefeller University, for their contributions to this work. We also would like to express our gratitude to the Animal Resources Department, and Environmental Protection Reserves & Wildlife Department, Ministry of Municipality and Environment in Qatar for their support and assistance for this project.

## Author contributions

Conceptualization: FA

Sampling: FA, IK

Methodology: FA, SR, QA, EJ, OF, GF, AT, KH, YS

Investigation: FA

Visualization: FA

Supervision: SR, QA, AAA

Writing—original draft: FA

Writing—review & editing: SR, QA, EJ, AAA

## Data and materials availability

All raw data of the reference genome assembly are available on VGP’s GenomeArk repository for raw data and assemblies (https://vgp.github.io/genomeark/Falco_rusticolus/). The resequencing data used in this study are available from Qatar Falcon Genome Project, but restrictions apply to the availability of these data, which were used under license for the current study, and so are not publicly available. The resequencing data are however available from the corresponding author (FA) upon request from bona fide researchers and with permission of Qatar Falcon Genome Project.

