## supplementary data for "Genomic, genetic and phylogenetic evidence for a new falcon species using chromosome-level genome assembly of the gyrfalcon and population genomics"

**This PDF file includes:**

Figs. S1 to S2
Tables S1 to S6

**Fig. S1.**
BUSCO genome-completeness analysis of the gyrfalcon reference genome

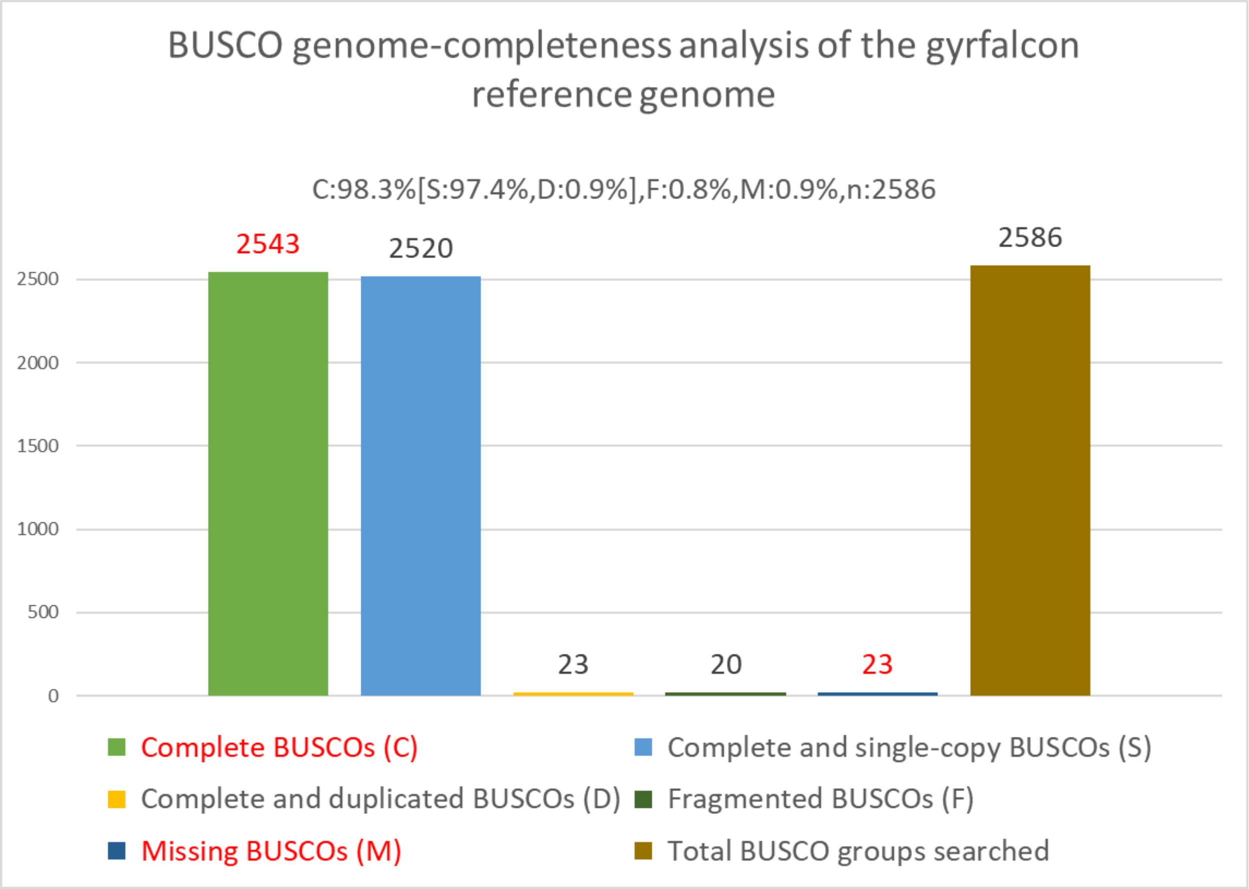

### Fig. S2.

Chronogram of falcon mitogenomes. Then nodes are labeled with the estimated age while the branch label shows the confidence level.

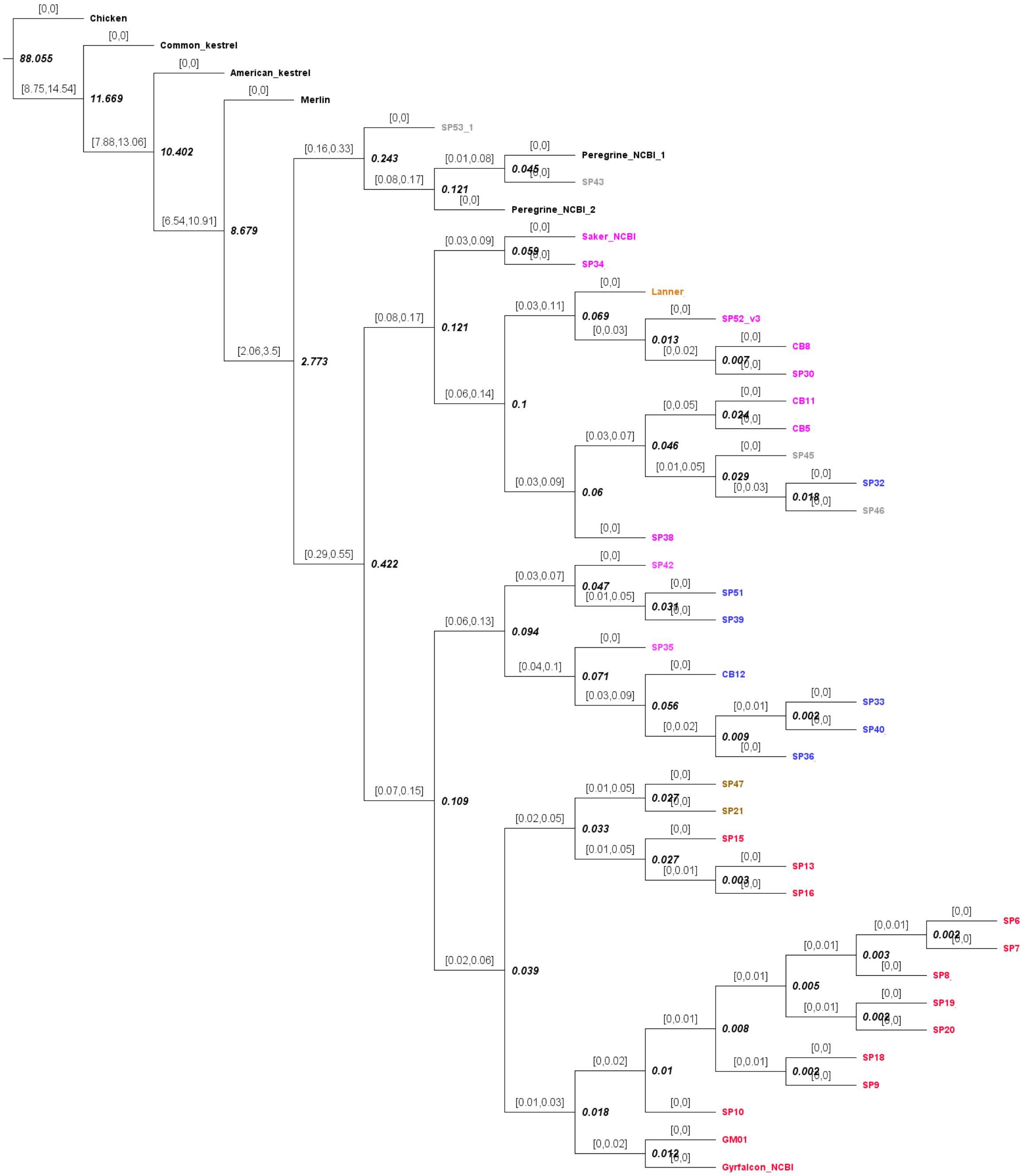

**Table S1. Primary and Alternative assembly stats**

Primary assembly stat

| | |
|---|---|
| Total sequence length | 1,195,847,496 |
| Total ungapped length | 1,190,501,682 |
| Gaps between scaffolds | 0 |
| Number of scaffolds | 133 |
| Scaffold N50 | 91,094,845 |
| Scaffold L50 | 6 |
| Number of contigs | 768 |
| Contig N50 | 15,323,486 |
| Contig L50 | 24 |
| Total number of chromosomes and plasmids | 25 |
| Number of component sequences (WGS or clone) | 133 |

Alternative assembly stats

| | |
|---|---|
| Total sequence length | 516,243,243 |
| Total ungapped length | 516,243,243 |
| Number of contigs | 8,370 |
| Contig N50 | 76,252 |
| Contig L50 | 2,120 |
| Total number of chromosomes and plasmids | 0 |
| Number of component sequences (WGS or clone) | 8,370 |

| Scaffold<br>All | Size (Gbp)<br>1,195,829,278 | Number<br>133 |
|---|---|---|
| Chromosome 1 | 126,922,963 | 1 |
| Chromosome 2 | 122,153,554 | 1 |
| Chromosome 3 | 121,276,792 | 1 |
| Chromosome 4 | 112,438,257 | 1 |
| Chromosome 5 | 92,439,089 | 1 |
| Chromosome 6 | 91,094,845 | 1 |

| | | |
|---|---|---|
| Chromosome 7 | 72,902,640 | 1 |
| Chromosome 8 | 65,417,759 | 1 |
| Chromosome 9 | 54,018,959 | 1 |
| Chromosome 10 | 38,161,925 | 1 |
| Chromosome 11 | 35,666,015 | 1 |
| Chromosome 12 | 33,884,028 | 1 |
| Chromosome 13 | 30,790,089 | 1 |
| Chromosome 14 | 24,097,451 | 1 |
| Chromosome 15 | 23,311,196 | 1 |
| Chromosome 16 | 8,171,160 | 1 |
| Chromosome 17 | 7,845,494 | 1 |
| Chromosome 18 | 6,718,660 | 1 |
| Chromosome 19 | 6,450,865 | 1 |
| Chromosome 20 | 6,217,219 | 1 |
| Chromosome 21 | 485,429 | 1 |
| Chromosome 22 | 425,072 | 1 |
| Chromosome W | 25,584,520 | 1 |
| Chromosome Z | 84,785,561 | 1 |
| unplaced | 4,569,736 | 108 |

**Table S2. Statistics of the three SNP panels developed using the falcon populations analyzed in this study**

| Panel Set | Total No. of raw variants (SNPs and indels) | Total No. of high-quality SNPs | Total No. of filtered SNPs (QC, pruned*) | No. Individuals used for analysis | Ts/Tv |
|---|---|---|---|---|---|
| Autosomal | 10,810,575 | 9,700,632 | 191,122* | 47 | 2.235 |
| Z | 1,011,596 | 861,171 | 4,975* | 37 (♀ only) | 2.135 |
| W | 62,742 | 47,735 | 60,441 (not pruned as it resemble a whole haplotype) | 37 (♀ only) | 2.391 |

**Table S3. Ancestry estimates of different falcon populations using ADMIXTURE, at K=2**

| Sample | Population | Gyrfalcon Ancestry Fraction ($Q_G$) | Saker Ancestry Fraction ($Q_S$) |
|---|---|---|---|
| All | Gyrfalcon (GF) (n=13) | 0.99999 | 0.00001 |
| All | Saker (SF) (n=8) | 0.00001 | 0.99999 |
| CB11 | Saker-like falcon (SLF) | 0.122571 | 0.877429 |
| SP42 | SLF | 0.399615 | 0.600385 |
| SP38 | SLF | 0.114843 | 0.885157 |
| CB08 | SLF | 0.187553 | 0.812447 |
| SP34 | SLF | 0.106561 | 0.893439 |
| SP35 | SLF | 0.105824 | 0.894176 |
| SP30 | SLF | 0.11 | 0.89 |
| SP52 | SLF | 0.149387 | 0.850613 |
| SP21 | Gyr-like falcon (GLF) | 0.935557 | 0.064443 |
| SP47 | GLF | 0.854768 | 0.145232 |
| SP50 | GLF | 0.507113 | 0.492887 |
| SP12 | Commercial hybrid (CH) | 0.793532 | 0.206468 |
| SP11 | CH | 0.61791 | 0.38209 |
| SP44 | CH | 0.627463 | 0.372537 |
| SP45 | CH | 0.818606 | 0.181394 |
| SP46 | CH | 0.938072 | 0.061928 |
| SP49 | CH | 0.350974 | 0.649026 |
| SP48 | CH | 0.9463 | 0.0537 |

**Table S4. Protein-coding genes annotated on the gyrfalcon's W chromosome (Accession Number NC_051209.1)**

| Gene | description | start position | end position | orientation | Exon count |
|---|---|---|---|---|---|
| *SYT4* | synaptotagmin-4-like | 15168729 | 15179015 | plus | 4 |
| *ZNF462* | zinc finger protein 462-like | 9852297 | 9859643 | plus | 2 |
| *DCAF12* | DDB1- and CUL4-associated factor 12-like | 8945048 | 8961020 | minus | 8 |
| *UBAP1* | ubiquitin-associated protein 1-like | 8783303 | 8795057 | plus | 6 |
| *LOC119140925* | uncharacterized LOC119140925 | 8234851 | 8239668 | plus | 3 |
| *DCC* | netrin receptor DCC-like | 7429787 | 7586607 | minus | 23 |
| *LAS2* | lung adenoma susceptibility protein 2-like | 7139486 | 7149893 | minus | 7 |
| *LOC119140922* | uncharacterized LOC119140922 | 6486359 | 6497667 | plus | 3 |
| *WDR7* | WD repeat-containing protein 7-like | 6361094 | 6475290 | minus | 21 |
| *NCBP1* | nuclear cap-binding protein subunit 1-like | 6091334 | 6155203 | plus | 15 |
| *NEK5* | NIMA related kinase 5 | 5862410 | 5892710 | plus | 20 |
| *LOC119140916* | uncharacterized LOC119140916 | 5705627 | 5728025 | plus | 5 |
| *SMRP1* | spermatid-specific manchette-related protein 1-like | 4654771 | 4662998 | plus | 7 |
| *LOC119140913* | uncharacterized LOC119140913 | 4263739 | 4268194 | plus | 4 |
| *VCP* | transitional endoplasmic reticulum ATPase-like | 3215897 | 3219246 | plus | 4 |
| *PSIP1* | lens epithelium-derived growth factor-like | 2765676 | 2772596 | plus | 5 |
| *ZNF131* | zinc finger protein 131-like | 2497552 | 2514541 | minus | 6 |
| *MALT1* | mucosa-associated lymphoid tissue lymphoma translocation protein 1-like | 1803423 | 1824102 | minus | 13 |
| *RIN2* | ras and Rab interactor 2-like | 1565489 | 1575435 | minus | 5 |
| *LOC119140901* | uncharacterized LOC119140901 | 75968 | 81107 | plus | 2 |
| *SLC1A3* | excitatory amino acid transporter 1-like | 3549063 | 3632341 | plus | 10 |

| | | | | | |
|---|---|---|---|---|---|
| *CKMT2* | creatine kinase S-type, mitochondrial-like | 10156326 | 10161157 | plus | 3 |
| *ERVV2* | endogenous retrovirus group V member 2 Env polyprotein-like | 15139217 | 15143289 | minus | 2 |
| *SMAD7* | mothers against decapentaplegic homolog 7-like | 1346117 | 1372897 | minus | 5 |
| *DAB2* | disabled homolog 2-like | 2528204 | 2552797 | plus | 13 |
| *CTIF* | CBP80/20-dependent translation initiation factor-like | 1177267 | 1325110 | plus | 13 |
| *RAB27B* | ras-related protein Rab-27B-like | 7045827 | 7075782 | minus | 6 |
| *CDC42SE2* | CDC42 small effector protein 2 | 3113656 | 3163753 | minus | 4 |
| *UBAP1* | ubiquitin-associated protein 1-like | 8831875 | 8866113 | minus | 12 |
| *RIT2* | GTP-binding protein Rit2-like | 15330898 | 15348542 | plus | 2 |
| *PCGF3* | polycomb group RING finger protein 3-like | 25202254 | 25322075 | plus | 13 |
| *FAM219A* | protein FAM219A-like | 4626305 | 4649337 | plus | 6 |
| *ISCA1* | iron-sulfur cluster assembly 1 homolog, mitochondrial-like | 4475863 | 4485297 | minus | 5 |
| *ISCA1* | iron-sulfur cluster assembly 1 homolog, mitochondrial-like | 4792570 | 4801709 | plus | 4 |
| *LOC119140868* | uncharacterized LOC119140868 | 5069273 | 5086188 | minus | 2 |
| *RASA1* | ras GTPase-activating protein 1-like | 4371881 | 4458936 | minus | 26 |
| *LOC119140866* | uncharacterized LOC119140866 | 7116423 | 7132830 | minus | 9 |
| *CCDC68* | coiled-coil domain-containing protein 68-like | 7042404 | 7045757 | plus | 5 |
| *CCDC68* | coiled-coil domain-containing protein 68-like | 7034073 | 7041156 | plus | 4 |
| *TCF4* | transcription factor 4-like | 6733431 | 6967265 | plus | 24 |
| *WASHC3* | WASH complex subunit 3-like | 23430893 | 23448704 | minus | 2 |

| | | | | | |
|---|---|---|---|---|---|
| *MIER3* | mesoderm induction early response protein 3-like | 23712425 | 23737080 | minus | 13 |
| *APBB1* | amyloid-beta A4 precursor protein-binding family A member 1-like | 23067889 | 23210030 | minus | 12 |
| *JMY* | junction-mediating and -regulatory protein-like | 24116260 | 24239432 | plus | 10 |
| *MAP3K1* | mitogen-activated protein kinase kinase kinase 1-like | 23634357 | 23702245 | plus | 21 |
| *HOMER1* | homer protein homolog 1-like | 24264816 | 24387599 | minus | 10 |
| *CHDW* | chromodomain-helicase-DNA-binding protein 1-like | 5347664 | 5417691 | minus | 37 |
| *CKAP2* | cytoskeleton associated protein 2 | 5823424 | 5835755 | plus | 12 |
| *THSD1* | thrombospondin type-1 domain-containing protein 1-like | 5773807 | 5794205 | minus | 4 |
| *ATP7B* | copper-transporting ATPase 2-like | 5923528 | 5949857 | plus | 22 |
| *TMEM272* | transmembrane protein 272-like | 5977189 | 5977539 | plus | 1 |
| *HAUS6* | HAUS augmin-like complex subunit 6 | 5742009 | 5763556 | plus | 6 |
| *ERVW1* | syncytin-A-like | 5607057 | 5608417 | minus | 2 |
| *NCBP1* | nuclear cap-binding protein subunit 1-like | 6116453 | 6134873 | minus | 9 |
| *DHRS12?* | dehydrogenase/reductase SDR family member 12-like | 5989908 | 6028556 | plus | 7 |
| *VPS36* | vacuolar protein sorting 36 homolog | 5801174 | 5823449 | minus | 14 |
| *ALG11* | ALG11 alpha-1,2-mannosyltransferase | 5895943 | 5900991 | minus | 3 |
| *PLIN2* | perilipin-2-like | 5729517 | 5737144 | plus | 8 |
| *NEK3* | NIMA related kinase 3 | 5849025 | 5862380 | plus | 16 |
| *VCP* | transitional endoplasmic reticulum ATPase-like | 3241710 | 3276407 | plus | 17 |
| *RFX3* | transcription factor RFX3-like | 2868546 | 3041951 | minus | 17 |
| *DUT* | deoxyuridine 5'-triphosphate nucleotidohydrolase-like | 8187310 | 8188577 | plus | 2 |
| *MBD2* | methyl-CpG-binding domain protein 2-like | 7166754 | 7207260 | plus | 7 |

| | | | | | |
|---|---|---|---|---|---|
| *ELAC1* | zinc phosphodiesterase ELAC protein 1-like | 8320688 | 8326479 | minus | 5 |
| *SMAD4* | mothers against decapentaplegic homolog 4-like | 8282177 | 8311271 | minus | 12 |
| *ME2* | NAD-dependent malic enzyme, mitochondrial | 8326725 | 8369371 | minus | 17 |
| *MEX3C* | RNA-binding E3 ubiquitin-protein ligase MEX3C-like | 8258841 | 8277165 | plus | 2 |
| *NTRK2* | BDNF/NT-3 growth factors receptor-like | 8444926 | 8691156 | minus | 19 |
| *ARRDC3* | arrestin domain-containing protein 3-like | 3941806 | 3978050 | minus | 13 |
| *FAM169A* | soluble lamin-associated protein of 75 kDa-like | 1497718 | 1522026 | plus | 12 |
| *CPLX4* | complexin 4 | 1648739 | 1669116 | plus | 3 |
| *DOT1L* | histone-lysine N-methyltransferase, H3 lysine-79 specific-like | 2479095 | 2480950 | minus | 2 |
| *SPIN* | spindlin-Z | 2129594 | 2443635 | minus | 15 |
| *GRPR* | gastrin releasing peptide | 1685670 | 1692932 | minus | 3 |
| *SEC11C* | SEC11 homolog C, signal peptidase complex subunit | 1703060 | 1710175 | minus | 9 |
| *RX2* | retinal homeobox protein Rx2-like | 1675236 | 1677755 | plus | 3 |
| *GNAQ* | guanine nucleotide-binding protein G(q) subunit alpha-like | 1919496 | 2055243 | plus | 7 |
| *PAIP1* | polyadenylate-binding protein-interacting protein 1-like | 2440606 | 2465603 | minus | 12 |
| *SNX2* | sorting nexin-2-like | 2328112 | 2378271 | plus | 17 |
| *LMAN1* | protein ERGIC-53-like | 1591648 | 1630893 | plus | 16 |
| *ZNF532* | zinc finger protein 532 | 1728861 | 1778264 | minus | 11 |
| *NTRK2* | BDNF/NT-3 growth factors receptor-like | 8463851 | 8526645 | plus | 8 |
| *IER3IP1* | immediate early response 3-interacting protein 1-like | 528990 | 534318 | minus | 3 |
| *SKOR2* | SKI family transcriptional corepressor 2-like | 560838 | 578876 | minus | 5 |
| *RNASEH1* | ribonuclease H-like | 414279 | 415426 | plus | 1 |

| | | | | | |
|---|---|---|---|---|---|
| *LOC119140782* | uncharacterized LOC119140782 | 411240 | 413826 | plus | 2 |
| *SMAD2* | mothers against decapentaplegic homolog 2 | 863089 | 910588 | minus | 13 |
| *ZBTB7C* | zinc finger and BTB domain-containing protein 7C-like | 951144 | 960269 | minus | 2 |
| *CERT1* | ceramide transfer protein-like | 403187 | 502390 | minus | 21 |
| *HNRNPK* | heterogeneous nuclear ribonucleoprotein K | 8759030 | 8777095 | plus | 18 |
| *ZFAND5* | AN1-type zinc finger protein 5-like | 22931249 | 22956771 | minus | 7 |
| *CDC37L1* | hsp90 co-chaperone Cdc37-like 1 | 22776537 | 22798315 | plus | 8 |
| *CZH18orf32* | UPF0729 protein C18orf32 homolog | 9147196 | 9150084 | minus | 4 |
| *DCAF12* | DDB1- and CUL4-associated factor 12-like | 8871333 | 8902088 | plus | 4 |
| *ZNF462* | zinc finger protein 462-like | 9873206 | 9886383 | plus | 3 |
| *RPL17* | 60S ribosomal protein L17-like | 9150359 | 9154025 | minus | 8 |
| *DIRAS2* | GTP-binding protein Di-Ras2-like | 9657726 | 9662530 | plus | 1 |
| *UBE2R2* | ubiquitin-conjugating enzyme E2 R2-like | 9053799 | 9082310 | minus | 5 |
| *NFIL3* | nuclear factor interleukin-3-regulated protein-like | 9582760 | 9596576 | minus | 4 |
| *LIPG* | endothelial lipase-like | 9168335 | 9177730 | plus | 10 |
| *MAPK4* | mitogen-activated protein kinase 4-like | 9361207 | 9408284 | plus | 14 |
| *UBQLN1* | ubiquilin-1-like | 9420196 | 9455847 | plus | 13 |
| *UBAP2* | ubiquitin-associated protein 2-like | 8973346 | 9052486 | plus | 32 |
| *ZNF462* | zinc finger protein 462-like | 9884850 | 9982453 | minus | 12 |
| *ERBIN* | erbin-like | 24989366 | 25109107 | minus | 29 |
| *LOC119140745* | uncharacterized LOC119140745 | 25163955 | 25166400 | minus | 2 |
| *LOC119140744* | uncharacterized LOC119140744 | 25131729 | 25133312 | minus | 2 |
| *LOC119140743* | uncharacterized LOC119140743 | 24866818 | 24888567 | plus | 5 |

| | | | | | |
|---|---|---|---|---|---|
| *TENT2* | poly(A) RNA polymerase GLD2-like | 24453214 | 24496138 | minus | 15 |
| *LOC119140740* | uncharacterized LOC119140740 | 24403807 | 24407563 | minus | 2 |
| *SCAMP1* | secretory carrier-associated membrane protein 1-like | 23917587 | 23938131 | plus | 6 |
| *UHRF2* | E3 ubiquitin-protein ligase UHRF2-like | 22474558 | 22522049 | plus | 3 |
| *PMAIP1* | phorbol-12-myristate-13-acetate-induced protein 1-like | 20987810 | 20988317 | plus | 2 |
| *PMAIP1* | phorbol-12-myristate-13-acetate-induced protein 1-like | 20969957 | 20970464 | plus | 2 |
| *LOC119140731* | uncharacterized LOC119140731 | 20818343 | 20866274 | minus | 2 |
| *TLN1* | talin-1-like | 20688470 | 20696553 | plus | 8 |
| *PPWD1* | peptidylprolyl isomerase domain and WD repeat-containing protein 1-like | 20322564 | 20333138 | plus | 5 |
| *ADAMTS6* | A disintegrin and metalloproteinase with thrombospondin motifs 6-like | 20138878 | 20284781 | minus | 23 |
| *LOC119140727* | uncharacterized LOC119140727 | 19696231 | 19702372 | plus | 4 |
| *ZSWIM6* | zinc finger SWIM domain-containing protein 6-like | 19586959 | 19659221 | plus | 6 |
| *LOC119140725* | uncharacterized LOC119140725 | 19308308 | 19313834 | minus | 2 |
| *DEPDC1B* | DEP domain-containing protein 1B-like | 19304134 | 19306422 | minus | 4 |
| *LOC119140722* | uncharacterized LOC119140722 | 19204329 | 19207579 | plus | 2 |
| *LOC119140721* | uncharacterized LOC119140721 | 19203096 | 19204355 | plus | 1 |
| *CELF4* | CUGBP Elav-like family member 4 | 17621302 | 18367914 | plus | 14 |
| *VLDLR* | very low-density lipoprotein receptor-like | 17181983 | 17182639 | minus | 1 |
| *LOC119140717* | protein NYNRIN-like | 15103084 | 15106209 | plus | 1 |
| *LOC119140716* | uncharacterized LOC119140716 | 15101851 | 15103178 | plus | 1 |
| *LOC119140714* | uncharacterized LOC119140714 | 15083291 | 15088830 | minus | 5 |

| | | | | | |
|---|---|---|---|---|---|
| *NYNRIN* | protein NYNRIN-like | 14203077 | 14205929 | plus | 1 |
| *LOC119140709* | uncharacterized LOC119140709 | 13438138 | 13440979 | plus | 4 |
| *HCN1* | potassium/sodium hyperpolarization-activated cyclic nucleotide-gated channel 1-like | 13289757 | 13370003 | plus | 5 |
| *HCN1* | potassium/sodium hyperpolarization-activated cyclic nucleotide-gated channel 1-like | 13173515 | 13211335 | plus | 3 |
| *ERVFRD-1* | syncytin-2-like | 12652166 | 12655653 | plus | 2 |
| *TNPO1* | transportin-1-like | 12150006 | 12207720 | minus | 24 |
| *LOC119140702* | uncharacterized LOC119140702 | 11759524 | 11762353 | minus | 2 |
| *HINT1* | histidine triad nucleotide-binding protein 1-like | 10939403 | 10943259 | minus | 4 |
| *SREK1* | splicing regulatory glutamine/lysine-rich protein 1-like | 24900574 | 24951448 | minus | 15 |
| *RNF38* | E3 ubiquitin-protein ligase RNF38-like | 25347291 | 25523437 | minus | 14 |
| *PIK3C3* | phosphatidylinositol 3-kinase catalytic subunit type 3 | 15671663 | 15754537 | minus | 27 |
| *SUB1* | activated RNA polymerase II transcriptional coactivator p15 | 21166913 | 21191255 | plus | 8 |
| *KCMF1* | E3 ubiquitin-protein ligase KCMF1-like | 22024860 | 22026686 | minus | 2 |
| *CENPK* | centromere protein K-like | 20293982 | 20306777 | minus | 5 |
| *APH1B* | gamma-secretase subunit Aph-1b-like | 20724935 | 20733788 | plus | 5 |
| *TMED7* | transmembrane emp24 domain-containing protein 7 | 21507704 | 21514944 | minus | 3 |
| *LOC119140670* | uncharacterized LOC119140670 | 21140118 | 21143029 | minus | 1 |
| *LOC119140668* | uncharacterized LOC119140668 | 20775435 | 20776330 | plus | 1 |
| *GOLPH3L* | Golgi phosphoprotein 3-like | 20913916 | 20938944 | plus | 3 |
| *TPM2* | tropomyosin beta chain-like | 20700465 | 20718868 | plus | 12 |
| *LOC119140664* | uncharacterized LOC119140664 | 21626630 | 21635184 | minus | 3 |

| | | | | | |
|---|---|---|---|---|---|
| *FEM1C* | protein fem-1 homolog C-like | 21304073 | 21327833 | plus | 3 |
| *HOOK3* | protein Hook homolog 3-like | 21865436 | 22020627 | plus | 24 |
| *FER* | tyrosine-protein kinase Fer-like | 21538765 | 21757517 | plus | 24 |
| *KCNN2* | small conductance calcium-activated potassium channel protein 2-like | 22151201 | 22267083 | plus | 9 |
| *ZFR* | zinc finger RNA-binding protein | 21020848 | 21067227 | minus | 22 |
| *NIPBL* | nipped-B-like protein | 20458703 | 20626320 | minus | 52 |
| *PLIN2* | perilipin-2-like | 4199867 | 4205288 | plus | 7 |
| *SMIM5* | small integral membrane protein 15-like | 19394602 | 19397141 | minus | 1 |
| *LOC119140653* | uncharacterized LOC119140653 | 20041919 | 20043389 | minus | 2 |
| *KIF2A* | kinesin-like protein KIF2A | 19816126 | 19909260 | minus | 22 |
| *LOC119140650* | uncharacterized LOC119140650 | 14220209 | 14222241 | minus | 2 |
| *LOC119140648* | uncharacterized LOC119140648 | 14214435 | 14225939 | minus | 2 |
| *LOC119140647* | uncharacterized LOC119140647 | 14371908 | 14374686 | plus | 2 |
| *SETBP1* | SET-binding protein-like | 14309296 | 14598732 | minus | 6 |
| *RICTOR* | rapamycin-insensitive companion of mTOR-like | 2557855 | 2667200 | plus | 41 |
| *APC* | adenomatous polyposis coli protein-like | 3423136 | 3457425 | minus | 8 |
| *APC* | adenomatous polyposis coli protein-like | 3334890 | 3414532 | minus | 14 |
| *ERVK-5* | endogenous retrovirus group K member 5 Gag polyprotein-like | 11426562 | 11427594 | minus | 2 |
| *ST8SIA3* | sia-alpha-2,3-Gal-beta-1,4-GlcNAc-R:alpha 2,8-sialyltransferase-like | 10974858 | 10979986 | plus | 4 |
| *LOC119140630* | uncharacterized LOC119140630 | 13724587 | 13725641 | plus | 1 |
| *FST* | follistatin-like | 13851941 | 13858457 | plus | 6 |
| *RNF165* | E3 ubiquitin-protein ligase RNF165-like | 11300833 | 11351843 | plus | 8 |
| *ERVFRD-1* | syncytin-2-like | 12326365 | 12333961 | minus | 4 |

| | | | | | |
|---|---|---|---|---|---|
| *LOC119140624* | uncharacterized protein C18orf25 | 11228908 | 11285149 | plus | 7 |
| *FECH* | ferrochelatase, mitochondrial-like | 11038157 | 11055901 | minus | 12 |
| *ONECUT2* | one cut domain family member 2-like | 10996193 | 11014353 | plus | 2 |
| *HMGCS1* | hydroxymethylglutaryl-CoA synthase, cytoplasmic-like | 13645007 | 13655530 | minus | 10 |
| *ZNF366* | zinc finger protein 366-like | 12021442 | 12033130 | minus | 4 |
| *LOC119140618* | uncharacterized LOC119140618 | 10844015 | 10846556 | plus | 1 |
| *SNX18* | sorting nexin-18-like | 10823373 | 10832963 | minus | 2 |
| *NARS1* | asparagine--tRNA ligase, cytoplasmic-like | 11060272 | 11074333 | minus | 16 |
| *PIAS2* | E3 SUMO-protein ligase PIAS2-like | 11523648 | 11556017 | minus | 14 |
| *ATP5F1A* | ATP synthase subunit alpha, mitochondrial-like | 11194979 | 11202762 | minus | 12 |
| *PAEP8* | protein mono-ADP-ribosyltransferase PARP8-like | 12910405 | 13061529 | minus | 26 |
| *NEDD4L* | E3 ubiquitin-protein ligase NEDD4-like | 10425380 | 10587533 | minus | 31 |
| *MAP1B* | microtubule-associated protein 1B-like | 11786893 | 11883416 | plus | 7 |

**Table S5. Number of W-linked genes involved in critical reproductive, physiological and pathological pathways using The PANTHER (Protein ANalysis THrough Evolutionary Relationships) Classification System**

| Pathway | Number of falcon W genes involved |
|---|---|
| Gonadotropin-releasing hormone receptor pathway (P06664) | 6 |
| Angiogenesis (P00005) | 5 |
| FGF signaling pathway (P00021) | 4 |
| TGF-beta signaling pathway (P00052) | 4 |
| EGF receptor signaling pathway (P00018) | 4 |
| Interleukin signaling pathway (P00036) | 3 |
| Insulin/IGF pathway-mitogen activated protein kinase kinase/MAP kinase cascade (P00032) | 3 |
| PDGF signaling pathway (P00047) | 3 |
| Wnt signaling pathway (P00057) | 2 |
| ATP synthesis (P02721) | 2 |
| Endothelin signaling pathway (P00019) | 2 |
| Metabotropic glutamate receptor group I pathway (P00041) | 2 |
| Metabotropic glutamate receptor group III pathway (P00039) | 1 |
| De novo pyrimidine deoxyribonucleotide biosynthesis (P02739) | 1 |
| Apoptosis signaling pathway (P00006) | 1 |
| JAK/STAT signaling pathway (P00038) | 1 |
| Ionotropic glutamate receptor pathway (P00037) | 1 |
| Alzheimer disease-presenilin pathway (P00004) | 1 |
| 5HT2 type receptor mediated signaling pathway (P04374) | 1 |
| Alzheimer disease-amyloid secretase pathway (P00003) | 1 |
| Interferon-gamma signaling pathway (P00035) | 1 |
| Integrin signalling pathway (P00034) | 1 |
| Inflammation mediated by chemokine and cytokine signaling pathway (P00031) | 1 |
| Heterotrimeric G-protein signaling pathway-Gq alpha and Go alpha mediated pathway (P00027) | 1 |
| Vasopressin synthesis (P04395) | 1 |
| Thyrotropin-releasing hormone receptor signaling pathway (P04394) | 1 |
| Toll receptor signaling pathway (P00054) | 1 |
| Ras Pathway (P04393) | 1 |
| T cell activation (P00053) | 1 |
| Oxytocin receptor mediated signaling pathway (P04391) | 1 |
| DNA replication (P00017) | 1 |
| PI3 kinase pathway (P00048) | 1 |
| Histamine H1 receptor mediated signaling pathway (P04385) | 1 |

| | |
|---|---|
| Cadherin signaling pathway (P00012) | 1 |
| Muscarinic acetylcholine receptor 1 and 3 signaling pathway (P00042) | 1 |
| Angiotensin II-stimulated signaling through G proteins and beta-arrestin (P05911) | 1 |
| Cortocotropin releasing factor receptor signaling pathway (P04380) | 1 |
| CCKR signaling map (P06959) | 1 |

**Table S6. Coding variants in W-linked genes in two falcon populations; peregrine (P) and Altai (A) along with their allele frequency. The impact variants were predicted using SnpEff; Red indicates High impact, and yellow indicated Moderate impact.**

| Gene | Variant 1 | Amino acid change | Coding variant | Transcript | Affected population | Variant 2 | Coding variant | Amino acid change | Transcript | Affected population |
|---|---|---|---|---|---|---|---|---|---|---|
| *NIPBL* | missense | p.Ala2012Thr | c.6034G>A | rna-XM_037372164.1 | P (3/3) | missense | c.3698T>C | p.Met1233Thr | rna-XM_037372164.1 | P (3/3) |
| *ADAMTS6* | missense | p.Ser877Pro | c.2629T>C | rna-XM_037372261.1 | A (10/10), P (3/3) | | | | | |
| *PIAS2* | missense | p.Ala7Val | c.20C>T | rna-XM_037372098.1 | A (10/10) | stop_lost | c.1720T>C | p.Ter574Argext*? | rna-XM_037372099.1 | P (3/3) |
| *NTRK2* | missense | p.Met48Ile | c.144G>C | rna-XM_037372386.1 | A (10/10) | missense | c.197T>C | p.Leu66Ser | rna-XM_037372386.1 | P (3/3) |
| *ELAC1* | missense | p.Gly170Asp | c.509G>A | rna-XM_037372458.1 | P (3/3) | | | | | |
| *MEX3C* | missense | p.Ser15Pro | c.43T>C | rna-XM_037372447.1 | A (10/10), P (3/3) | missense | c.234T>G | p.Asp78Glu | rna-XM_03737244 | P (3/3) |

| | | | | | | | | | | |
|---|---|---|---|---|---|---|---|---|---|---|
| | | | | | | | | | 7.1 | |
| *APBA1* | miss ense | p.Glu235 Ala | c.704 A>C | rna-XM_0 37372491. 1 | P (3/3) | | | | | |
| *CDC37L1* | miss ense | p.Leu315 Pro | c.944T >C | rna-XM_0 37372342. 1 | P (3/3) | | | | | |
| *NCBP1* | miss ense | p.Val388I le | c.1162 G>A | rna-XM_0 37372581. 1 | P (3/3) | | | | | |
| *DCAF12* | miss ense | p.Arg97G ln | c.290 G>A | rna-XM_0 37372590. 1 | P (3/3) | | | | | |
| *WDR7* | miss ense | p.Arg143 Gln | c.428 G>A | rna-XM_0 37372582. 1 | P (3/3) | | | | | |
| *NYNRIN-LI KE* | miss ense | p.Leu264 Pro | c.791T >C | rna-XM_0 37372583. 1 | A (10/10), P (3/3) | | | | | |
| *UPF0729* | miss ense | p.Lys71G lu | c.211 A>G | rna-XM_0 37372340. 1 | P (3/3) | | | | | |
| *ZNF462* | miss ense | p.Pro40S er | c.118C >T | rna-XM_0 37372337. 1 | A (10/10) | missense | c.79G>A | p.Val27Met | rna-X M_037 37233 | P (3/3) |

| | | | | | | | | | | |
|---|---|---|---|---|---|---|---|---|---|---|
| | | | | | | | | | 7.1 | |
| *PLIN2* | missense | p.Arg110Thr | c.327_329delTAGinsAAC | rna-XM_037372468.1 | A (10/10), P (3/3) | | | | | |
| *PLIN2* | missense | p.Thr348Met | c.1043C>T | rna-XM_037372153.1 | A (10/10), P (3/3) | stop_gained | c.1310T>A | p.Leu437* | rna-XM_037372153.1 | P (3/3) |
| *UBQLN1* | missense | p.Met178Val | c.532A>G | rna-XM_037372310.1 | P (3/3) | missense | c.788A>G | p.Asn263Ser | rna-XM_037372310.1 | A (10/10) |
| *MAPK4* | missense | p.Asn498Ser | c.1493A>G | rna-XM_037372315.1 | P (3/3) | | | | | |
| *APC* | missense | p.Thr123Met | c.368C>T | rna-XM_037372129.1 | A (10/10), P (3/3) | | | | | |
| *ISCA1* | missense | p.Asn169Ser | c.506A>G | rna-XM_037372523.1 | P (3/3) | frameshift | c.28_29insG | p.Arg12fs | rna-XM_037372524.1 | P (3/3) |
| *LOC119140868* | missense | p.Ile205Thr | c.614T>C | rna-XM_037372520.1 | A (8/10) | missense | c.224C>T | p.Pro75Leu | rna-XM_037372520.1 | P (3/3) |

| | | | | | | | | | | |
|---|---|---|---|---|---|---|---|---|---|---|
| *SMRP1* | missense | p.Ser98Asn | c.293G>A | rna-XM_037372577.1 | A (10/10), P (3/3) | | | | | |
| *LOC119140916* | missense | p.Cys41Arg | c.121T>C | rna-XM_037372579.1 | A (9/10) | missense | c.712C>T | p.Arg238Trp | rna-XM_037372579.1 | P (3/3) |
| *RASA1* | conservative_inframe_insertion | p.Gly90_Gln91insGlnGly | c.268_269insGCCAGG | rna-XM_037372517.1 | A (10/10), P (3/3) | | | | | |
| *CTIF* | missense | p.Gln347Pro | c.1040A>C | rna-XM_037372552.1 | P (3/3) | | | | | |
| *SMAD7* | missense | p.Ser50Gly | c.148A>G | rna-XM_037372560.1 | P (3/3) | | | | | |
| *LMAN1* | missense | p.Leu15Phe | c.43C>T | rna-XM_037372395.1 | P (3/3) | splice_acceptor_variant&intron_variant | c.481-3_481-2insT | | rna-XM_037372392.1 | P (3/3) |
| *RAX* | missense | p.Ile82Val | c.243_244delGAins | rna-XM_037372406.1 | P (3/3) | | | | | |

| | | | | | | | | | | |
|---|---|---|---|---|---|---|---|---|---|---|
| | | | AG | | | | | | | |
| *GRP* | missense | p.Ser239Pro | c.711_715delACTTTinsGCTTC | rna-XM_037372414.1 | A (10/10), P (3/3) | | | | | |
| *RNHL* | missense | p.Gln54Arg | c.161A>G | rna-XM_037372381.1 | P (3/3) | stop_gained | c.547C>T | p.Arg183* | rna-XM_037372381.1 | A (10/10) |
| *ZBTB7C* | missense | p.Ile596Met | c.1788A>G | rna-XM_037372375.1 | P (3/3) | missense | c.1594A>C | p.Lys532Gln | rna-XM_037372375.1 | P (3/3) |
| *MALT1* | missense | p.Gly567Ser | c.1699G>A | rna-XM_037372568.1 | P (3/3) | missense | c.1478G>A | p.Cys493Tyr | rna-XM_037372568.1 | P (3/3) |
| *PAIP1* | missense | p.Asn425Ile | c.1274A>T | rna-XM_037372403.1 | P (3/3) | | | | | |
| *DOT1L* | missense | p.Ser106Thr | c.316T>A | rna-XM_037372433.1 | A (10/10), P (3/3) | | | | | |
| *ZNF131* | missense | p.Ala19Ser | c.54_55delGGinsT | rna-XM_037372570.1 | P (3/3) | | | | | |

| | | | | | | | | | | |
|---|---|---|---|---|---|---|---|---|---|---|
| | | | T | | | | | | | |
| *DAB2* | missense | p.Leu225Val | c.673C>G | rna-XM_037372555.1 | P (3/3) | | | | | |
| *RICTOR* | missense | p.His1548Arg | c.4643A>G | rna-XM_037372135.1 | A (5/10) | | | | | |
| *ATP7B* | missense&splice_region_variant | p.Gln1290Arg | c.3869A>G | rna-XM_037372478.1 | P (3/3) | | | | | |
| *CCDC68* | missense | p.His49Arg | c.146A>G | rna-XM_037372515.1 | P (3/3) | | | | | |
| *DYNAP* | missense | p.Gly373Arg | c.1117G>A | rna-XM_037372516.1 | A (10/10) | missense | c.986T>A | p.Ile329Asn | rna-XM_037372516.1 | A (10/10) |
| *LAS2* | missense | p.Tyr206Ser | c.617A>C | rna-XM_037372584.1 | P (3/3) | missense | c.223T>C | p.Cys75Arg | rna-XM_037372584.1 | P (3/3) |

| | | | | | | | | | | |
|---|---|---|---|---|---|---|---|---|---|---|
| *DCC* | missense | p.Thr426Met | c.1277C>T | rna-XM_037372586.1 | A (10/10) | missense | c.2132A>G | p.Asp711Gly | rna-XM_037372586.1 | P (3/3) |
| *DUT* | missense | p.Val83Ile | c.247G>A | rna-XM_037372460.1 | P (3/3) | | | | | |
| *UBAP2* | missense | p.Arg169His | c.506G>A | rna-XM_037372295.1 | A (10/10) | splice_acceptor_variant&splice_region_variant&intron_variant | c.973-21_973-1delCTTTTTTTTTTCTTTTGTTAGinsTTTTT TTTTTTTATTTTGTTAG | | rna-XM_037372304.1 | P (3/3) |
| *LIPG* | missense | p.Thr348Met | c.1043C>T | rna-XM_037372326.1 | A (10/10), P (3/3) | missense | c.1061A>G | p.Asn354Ser | rna-XM_037372326.1 | A (10/10), P (3/3) |
| *ONECUT2* | missense | p.Thr263Pro | c.787A>C | rna-XM_037372108.1 | P (3/3) | missense | c.1378T>C | p.Ser460Pro | rna-XM_037372108.1 | P (3/3) |

| | | | | | | | | | | |
|---|---|---|---|---|---|---|---|---|---|---|
| *NARS1* | missense&splice_region_variant | p.Asp15Glu | c.45T>G | rna-XM_037372100.1 | P (3/3) | | | | | |
| *18ORF25 HOMOLOG* | missense | p.Leu55Val | c.163C>G | rna-XM_037372112.1 | P (3/3) | missense | c.391T>C | p.Ser131Pro | rna-XM_037372112.1 | P (3/3) |
| *LOC119140702* | missense | p.Ala249Asp | c.746C>A | rna-XM_037372237.1 | P (3/3) | frameshift | c.983_984insAACA | p.Pro330fs | rna-XM_037372237.1 | P (3/3) |
| *MAP1B* | missense | p.Pro1332Ser | c.3994C>T | rna-XM_037372093.1 | P (3/3) | missense | c.4213T>C | p.Cys1405Arg | rna-XM_037372093.1 | P (3/3) |
| *ERVFRD-1* | missense | p.Thr123Met | c.368C>T | rna-XM_037372118.1 | P (3/3) | missense | c.163A>G | p.Lys55Glu | rna-XM_037372116.1 | P (3/3) |
| *PARP8* | missense | p.Arg266His | c.797G>A | rna-XM_037372096.1 | P (3/3) | | | | | |

| | | | | | | | | | | |
|---|---|---|---|---|---|---|---|---|---|---|
| *HCN1* | missense | p.Ser175Asn | c.524G>A | rna-XM_037372243.1 | P (3/3) | | | | | |
| *ERVK-18* | missense | p.Arg222Gln | c.665G>A | rna-XM_037372244.1 | P (3/3) | missense | c.932C>T | p.Thr311Met | rna-XM_037372244.1 | P (3/3) |
| *HMGCS1* | missense | p.Arg75Lys | c.224G>A | rna-XM_037372107.1 | P (3/3) | missense&splice_region_variant | c.16A>G | p.Ile6Val | rna-XM_037372107.1 | P (3/3) |
| *NYNRIN* | missense | p.Lys845Arg | c.2534A>G | rna-XM_037372247.1 | A (10/10), P (3/3) | missense | c.2651G>A | p.Arg884His | rna-XM_037372247.1 | P (3/3) |
| *LOC119140648* | missense | p.Ile32Leu | c.94A>C | rna-XM_037372145.1 | P (3/3) | stop_gained | c.376C>T | p.Arg126* | rna-XM_037372145.1 | P (3/3) |
| *VLDLR* | missense | p.Glu8Asp | c.24A>T | rna-XM_037372252.1 | P (3/3) | | | | | |
| *CELF4* | missense | p.Ala271Thr | c.811G>A | rna-XM_037372253.1 | P (3/3) | | | | | |
| *NYNRIN-LIKE* | missense | p.Met947Ile | c.2841G>A | rna-XM_037372255.1 | P (3/3) | missense | c.3038G>A | p.Arg1013Lys | rna-XM_03737225 | A (10/10), P (3/3) |

| | | | | | | | | | | |
|---|---|---|---|---|---|---|---|---|---|---|
| | | | | | | | | | 5.1 | |
| *NYNRIN-LIKE* | missense | p.Thr309Ala | c.925A>G | rna-XM_037372257.1 | P (3/3) | | | | | |
| *ZSWIM6* | missense&conservative_inframe_deletion | p.Ala47_Ala52delinsProProAlaAlaVal | c.139_155delGCCGCCGCCGCCGTCGCinsCCCCCCGCCGCCGT | rna-XM_037372258.1 | P (3/3) | | | | | |
| *NYNRIN-LIKE* | missense | p.Lys18Glu | c.52A>G | rna-XM_037372260.1 | P (3/3) | missense | c.2425T>C | p.Cys809Arg | rna-XM_037372260.1 | P (3/3) |
| *KIF2A* | missense | p.Pro700Leu | c.2099C>T | rna-XM_037372150.1 | P (3/3) | | | | | |
| *PPWD1* | missense | p.Val127Met | c.379G>A | rna-XM_037372263.1 | P (3/3) | | | | | |
| *APH1B* | missense | p.Ala10Thr | c.28G>A | rna-XM_037372210.1 | P (3/3) | | | | | |

| | | | | | | | | | | |
|---|---|---|---|---|---|---|---|---|---|---|
| *NYNRIN-LIKE* | missense | p.Pro76Ala | c.222_226delACGACnsGCGAG | rna-XM_037372206.1 | P (3/3) | missense | c.244G>A | p.Glu82Lys | rna-XM_037372206.1 | P (3/3) |
| *LOC119140731* | missense | p.Pro609Leu | c.1826C>T | rna-XM_037372265.1 | A (10/10) | missense | c.488A>G | p.His163Arg | rna-XM_037372265.1 | P (3/3) |
| *ZFR* | missense | p.Ala1055Gly | c.3164C>G | rna-XM_037372166.1 | P (3/3) | | | | | |
| *NYNRIN-LIKE* | missense | p.Val164Ala | c.491T>C | rna-XM_037372207.1 | P (3/3) | | | | | |
| *FER* | missense | p.Met316Thr | c.947T>C | rna-XM_037372176.1 | P (3/3) | missense | c.2489G>A | p.Arg830Lys | rna-XM_037372176.1 | P (3/3) |
| *HOOK3* | missense | p.Thr191Met | c.572C>T | rna-XM_037372184.1 | P (3/3) | | | | | |
| *WASHC3* | missense | p.Tyr12Cys | c.35A>G | rna-XM_037372496.1 | A (10/10), P (3/3) | missense | c.89C>T | p.Ala30Val | rna-XM_037372496.1 | P (3/3) |
| *MAP3K1* | missense | p.Leu109Pro | c.326T>C | rna-XM_037372487. | P (3/3) | missense&splice_ | c.1256T>C | p.Met419Thr | rna-XM_037 | P (3/3) |

| | | | | | | | | | | |
|---|---|---|---|---|---|---|---|---|---|---|
| | | | | 1 | | region_variant | | | 372487.1 | |
| *MIER3* | missense | p.Arg512His | c.1535G>A | rna-XM_037372493.1 | P (3/3) | | | | | |
| *JMY* | missense | p.His836Arg | c.2507A>G | rna-XM_037372488.1 | P (3/3) | | | | | |
| *NYNRIN-LIKE* | missense | p.Arg613Gly | c.1837A>G | rna-XM_037372273.1 | A (10/10), P (3/3) | missense | c.1879A>G | p.Thr627Ala | rna-XM_037372273.1 | P (3/3) |
| *TENT2* | missense | p.Arg367Ser | c.1099C>A | rna-XM_037372274.1 | P (3/3) | missense | c.482G>A | p.Arg161His | rna-XM_037372274.1 | P (3/3) |
| *LOC119140743* | conservative_inframe_deletion | p.Leu22del | c.64_66delCTC | rna-XM_037372275.1 | P (3/3) | | | | | |
| *LOC119140744* | missense | p.Arg33Leu | c.98G>T | rna-XM_037372277.1 | P (3/3) | missense | c.239C>T | p.Pro80Leu | rna-XM_037372277.1 | P (3/3) |
| *NYNRIN-LI* | miss | p.Pro174 | c.520 | rna-XM_0 | A | frameshi | c.176del | p.Arg59fs | rna-X | P (3/3) |

| | | | | | | | | | |
|---|---|---|---|---|---|---|---|---|---|
| *KE* | ense | Thr | C>A | 37372278.1 | (10/10) | ft | G | | M_037372277.1 |
| *RNF38* | missense | p.Tyr59Cys | c.176A>G | rna-XM_037372225.1 | P (3/3) | | | | |
| *CHD1* | frameshift&splice_donor_variant&splice_region_variant&synonymous_variant&intron_variant | p.Gln771fs | c.2312_2313+26delAGGTAATTTATTTATTTTTTTTTTTTTTinsAGGTAATTTATTTTTTTTTTTTTTTAA | rna-XM_037372482.1 | A (10/10), P (3/3) | | | | |
| DCAF1 | frameshift | p.Ser198fs | c.591_592insA | rna-XM_037372590.1 | P (3/3) | | | | |
| THSD1 | stop_lost | p.Ter495Argext*? | c.1483T>C | rna-XM_037372479. | P (3/3) | | | | |

| | | | | | |
|---|---|---|---|---|---|
| | | | | 1 | |
| SPINW | stop_gained | p.Arg10* | c.28C>T | rna-XM_037372415.1 | P (3/3) |
| NEK3 | splice_acceptor_variant&intron_variant | | c.805-2A>G | rna_XM_037372467.1-1 | P (3/3) |
| HINT1 | frameshift | p.Gly85fs | c.240_241insGCCCG | rna-XM_037372236.1 | P (3/3) |
| ARRDC3 | frameshift | p.Val99fs | c.282_283insT | rna-XM_037372443.1 | P (3/3) |
| LOC105374013 | stop_gained | p.Arg128* | c.382C>T | rna-XM_037372127.1 | P (3/3) |
